# Target RNA abundance controls the collateral activity of RfxCas13d in human cells and zebrafish embryos

**DOI:** 10.64898/2026.04.07.716849

**Authors:** Honglin Chen, Wenxin Hu, Valeria Impicciche, Gurjeet Jagjeet Singh, Joshua King, Carolyn Shembrey, Priyank Rawat, Joshua ML Casan, Srdjan Boskovic, Scott Paterson, Wei Zhao, Sharon R Lewin, Ricky W Johnstone, Benjamin M Hogan, Stephin J Vervoort, Joseph A Trapani, Kazuhide S Okuda, Mohamed Fareh

## Abstract

Collateral RNA cleavage by CRISPR-Cas13 effectors presents a critical obstacle to their application in biological research and therapeutics, yet the molecular determinants of this activity remain poorly understood. Here, we systematically investigate the collateral activity of various Cas13 variants in human cells in vitro and zebrafish embryos in vivo. Among these nucleases, *Rfx*Cas13d displays robust collateral activity that is highly dependent on target abundance. Targeting moderately expressed RNAs activates only a limited subset of RfxCas13d molecules, resulting in selective degradation of ectopically expressed transcripts, while endogenous RNAs remain largely protected. This selectivity indicates higher accessibility to exogenous RNAs through spatial proximity and temporal colocalization with activated RfxCas13d nuclease domains. In contrast, the recognition of highly abundant RNA targets drives simultaneous activation of a large fraction of cellular RfxCas13d, leading to widespread collateral cleavage of cellular RNAs, disruption of proteome homeostasis, and consequent cell toxicity and developmental defects in zebrafish embryos. Notably, transgenic zebrafish with target RNA expression restricted to endothelial or neuronal cell lineages exhibit localized collateral activity, leading to tissue-specific developmental abnormalities and motility deficits. These findings reveal that *Rfx*Cas13d’s collateral activity is threshold-dependent, with abundant target RNA acting as a molecular switch for widespread collateral RNA degradation. This work underscores the need for careful consideration of target abundance when deploying *Rfx*Cas13d, and highlights *Psp*Cas13b as an alternative for RNA silencing free of collateral activity.

## INTRODUCTION

The discovery of CRISPR (Clustered Regularly Interspaced Short Palindromic Repeats) and its associated Cas (CRISPR-associated) proteins has unlocked unprecedented opportunities for precise genome and transcriptome editing [1, 2]. Among these remarkable CRISPR tools, Cas13 enzymes enable exclusive recognition and cleavage of single-stranded RNA without altering genomic DNA [2–4]. Numerous reports have leveraged Cas13 for various RNA editing applications including silencing of coding and non-coding cellular RNA, suppression of pathogenic viruses, and precise RNA base-editing [5–9].

Cas13 uses a single CRISPR RNA (crRNA) that base-pairs with the target RNA, triggering conformational changes in the Cas13 protein and activating its two HEPN (higher eukaryotes and prokaryotes nucleotide-binding) RNase domains, which then mediate potent RNA cleavage [2, 3, 10–12]. In addition, Cas13’s spacer sequence, which is approximately 22–30 nucleotides long, coupled with its low mismatch tolerance, ensures high targeting fidelity and minimizes off-target effects [2, 5, 13]. This distinguishes Cas13 from eukaryotic RNA interference machinery and the DNA-targeting Cas9 enzyme, both of which recognise shorter target sequences [14–18]. This long target recognition motif makes Cas13 ideal for programmable and highly specific RNA silencing for research and therapeutic applications [19–21]. However, Cas13 targeting specificity can be compromised by collateral activity, where target-activated HEPN nuclease domains trigger indiscriminate degradation of nearby RNA molecules. Mechanistically, the conformational changes following specific spacer-target base-pairing expose an activated HEPN domain on Cas13’s surface, leading to nonspecific degradation of proximal RNA [10, 22, 23]. In bacterial systems, Cas13-mediated collateral RNA degradation is believed to trigger cell dormancy or suicide, serving as an additional altruistic defence mechanism against phage infection [2, 10, 22, 24]. Interestingly, this collateral activity has been repurposed *in vitro* for highly sensitive RNA diagnostics [2, 25, 26]. However, such assays typically use RNA concentrations far exceeding physiological levels, and the free diffusion of RNA in these artificial systems increases the likelihood of Cas13-RNA interactions, thereby amplifying collateral effects. By contrast, RNA expression, abundance, and localisation are tightly regulated in eukaryotic cells, potentially limiting RNA interaction with the HEPN nuclease domains and modulating their susceptibility to collateral degradation [27, 28]. Despite this, the true extent of Cas13 collateral activity in eukaryotic systems remains poorly defined, with existing studies reporting conflicting findings [2, 29–41]. This inconsistency underscores a critical knowledge gap that must be addressed to safely and effectively deploy Cas13 technologies in research and therapeutic applications.

Here, we performed a systematic head-to-head comparison of the on target and collateral activity of four Cas13 variants in human cells. Fluorescence, proteomic and transcriptomic analysis demonstrated that *Lwa*Cas13a and *Psp*Cas13b possess robust on-target activity with minimal off-target effects, whereas the high fidelity *hf*Cas13d suffers from an impaired on-target cleavage efficacy. In contrast, RfxCas13d exhibited unique, RNA threshold-dependent collateral activity. Specifically, targeting moderately expressed transcripts triggered selective degradation of exogenous RNAs, whereas endogenous transcripts were largely spared. Conversely, targeting of high-abundance RNAs induced global collateral RNA degradation and deregulation of the proteome, resulting in cellular toxicity and developmental defects in zebrafish embryos. These findings reveal the tunability of *Rfx*Cas13d collateral activity in human cells and zebrafish embryos, with target abundance acting as a molecular switch.

## RESULTS

### Cas13 collateral activity varies between orthologs

The poor understanding of the molecular basis underlying Cas13 collateral activity undermines the utility of this family of programmable RNA nucleases. To address this, we investigated the collateral activity of three Cas13 orthologs in human cells. We first used a fluorescence–based assay to examine the on-target and collateral activity against the target mCherry (red) and non-target EGFP (green) transcripts. In this assay, Cas13 enzymes are programmed with two different crRNAs to specifically silence mCherry mRNA in HEK293T cells, whereas the nontarget EGFP mRNA is used to evaluate Cas13’s collateral activity (**Figure 1a, Suppl. Figure 1a**).

**Figure 1.**
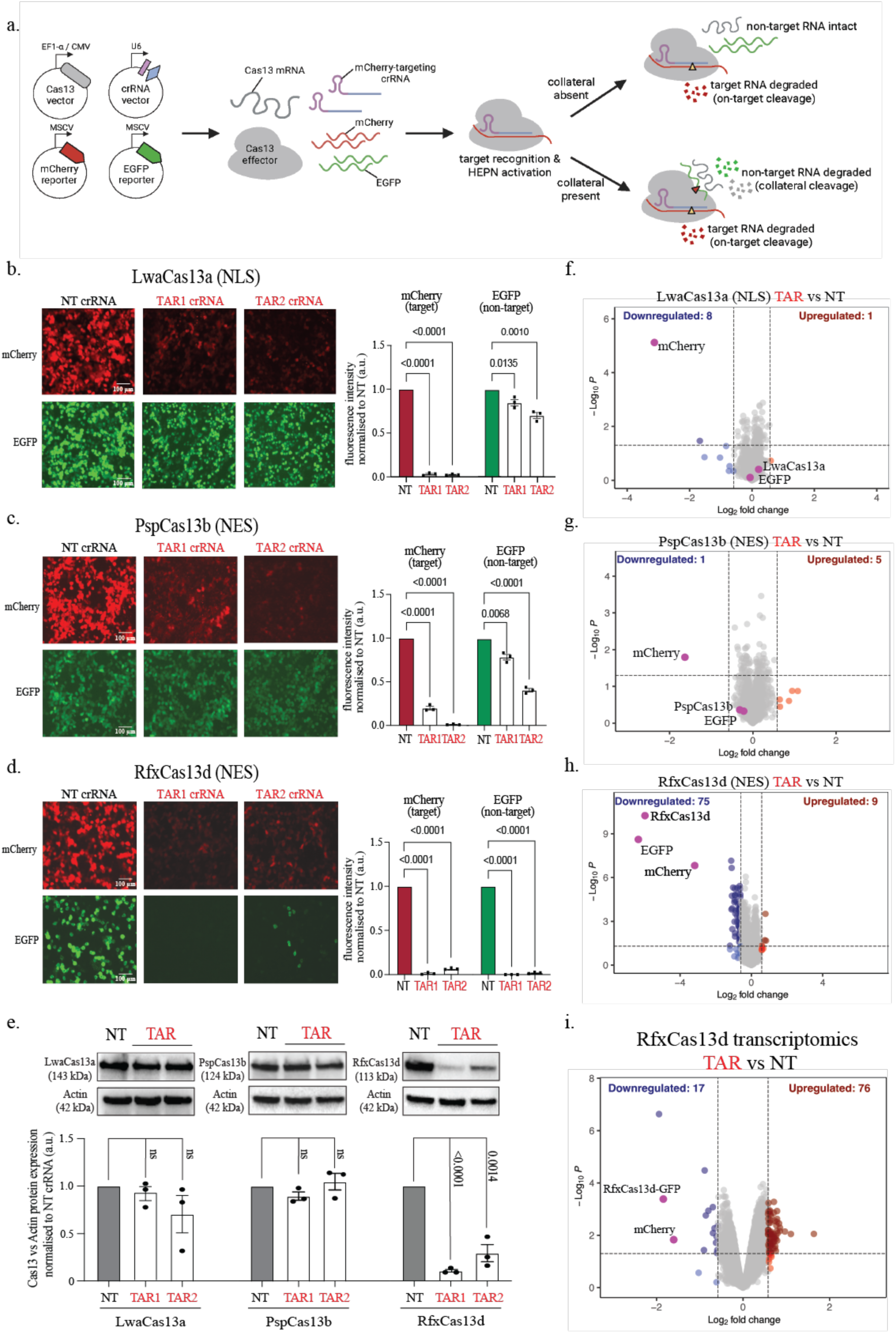
**a,** Schematic of CRISPR/Cas13 plasmid transfection dual fluorescence mCherry (target) and EGFP (non-target) silencing assay to assess three Cas13 orthologues (LwaCas13a, PspCas13b, RfxCas13d) on-target and collateral cleavage. **b-d, Left,** Representative fluorescence microscopy images of the silencing of mCherry and EGFP transcripts by three Cas13 orthologs with two potent mCherry-targeting crRNAs (TAR1, TAR2) versus a non-targeting (NT) control crRNA in HEK293T cells (n = 3); **Right,** Quantification of Cas13 silencing of mCherry and EGFP transcripts with either NT or mCherry-targeting crRNAs. Data points in the graph are averages of the normalized mean fluorescence from four to six representative fields of view imaged (n = 3). The data are represented in arbitrary units (a.u.). Errors are the s.e.m. and p values of Student’s t-test are indicated (95% confidence interval). **e, Top,** Representative Western blot of each Cas13 ortholog effector protein levels and β actin protein levels in HEK293T cells expressing either mCherry-targeting (TAR) crRNAs or NT crRNA. **Bottom,** Quantification of Cas13 protein silencing normalised to βactin protein levels (n = 3). The data are represented in arbitrary units (a.u.). Errors are the s.e.m. and p values of Student’s t-test are indicated (95% confidence interval). **f-h,** Volcano plots showing the proteome of HEK293T cells expressing mCherry, EGFP, mCherry-targeting or NT crRNA paired with different Cas13 orthologs. Data points represent proteins, with log_2_FC > 0.58 (upregulation, blue) or log_2_FC < −0.58 (downregulation, red) and p < 0.05 indicating significantly differential expression. Total downregulated and upregulated proteins are shown at the top corners. Exogenous Cas13, mCherry and EGFP are labelled purple. Results were analysed by an unpaired two-tailed Student’s t-test (n = 5). **i,** Volcano plots showing the transcriptome of HEK293T cells expressing mCherry, EGFP, mCherry-targeting or NT crRNA paired with different Cas13 orthologs. Data points represent RNAs, with log_2_FC > 1 (upregulation, blue) or log_2_FC < −1 (downregulation, red) and p < 0.05 indicating significantly differential expression. Results were analysed by an unpaired two-tailed Student’s t-test (n = 5).

*Lwa*Cas13a and *Psp*Cas13b orthologs exhibited potent on-target activity that led to 95-99% reduction in mCherry fluorescence, while EGFP fluorescence was reduced by only 15-25% (**Figure 1b, 1c**). *Rfx*Cas13d also showed a potent on-target activity that mediated ∼99% silencing of mCherry (**Figure 1d**). However, target recognition unleashed a substantial collateral activity of *Rfx*Cas13d that also suppressed ∼99% of the non-target EGFP fluorescence (**Figure 1d**). This was also confirmed by flow cytometry, where *Rfx*Cas13d exhibited a significant reduction of mCherry and EGFP double-positive cells and increased double-negative cells. In contrast, *Psp*Cas13b downregulated the expression of the target mCherry only (**Suppl. Figure 1b)**. We also questioned whether cellular localisation of *Rfx*Cas13d can affect the level of its collateral activity. The data revealed that *Rfx*Cas13d’s collateral activity remained potent regardless of its nuclear or cytoplasmic localisation, and that *Lwa*Cas13a and *Psp*Cas13b didn’t present collateral cleavage regardless of their localisation (**Figure 1d, Suppl. Figure 2b**). Using sequence alignment, we confirmed poor sequence complementarity between EGFP mRNA and mCherry crRNAs, indicating that the loss of EGFP fluorescence is due to the collateral activity of RfxCas13d activated by the recognition of mCherry target transcripts. We further confirmed that in the absence of the mCherry target, the mCherry-specific crRNA is unable to silence EGFP fluorescence (**Suppl. Figure 2c**).

This demonstrates that the loss of EGFP fluorescence is due to the collateral activity upon target recognition and *Rfx*Cas13d nuclease domain activation rather than direct targeting of EGFP by mismatched mCherry-targeting crRNAs. Western blot analysis confirmed that the collateral activity of *Rfx*Cas13d led to the cleavage of its own RNA and subsequent protein downregulation (**Figure 1e**). In contrast, target-activated *Lwa*Cas13a and *Psp*Cas13b orthologs remained free of any collateral activity against their own Cas13 transcripts (**Figure 1e**). In addition, the collateral activity of *Rfx*Cas13d led to a noticeable cleavage of the ribosomal RNA (rRNA) 28S, whereas target-activated *Psp*Cas13b did not show any collateral cleavage against 28S rRNA (**Suppl. Figure 2d**). *Rfx*Cas13d also exhibited similar properties of collateral activity when programmed to target another exogenous transcript (Tat HIV RNA) ectopically expressed in HEK293T cells (**Suppl. Figure 3**). This suggested that the collateral activity against ectopically expressed transcripts is independent of target identity or cellular localisation of *Rfx*Cas13d. Collectively, these findings indicated that, unlike *Lwa*Cas13a and *PspCas13b*, *Rfx*Cas13d ortholog possesses a substantial target-dependent collateral activity that can efficiently degrade ectopically expressed non-target RNA.

### The high-fidelity Cas13d (*hf*Cas13d) exhibits altered on-target activity

Tong et al. [34] engineered a high-fidelity variant of *Rfx*Cas13d (*hf*Cas13d) through targeted mutagenesis, which abolished its collateral activity while retaining on-target cleavage. Using the same RfxCas13d crRNA targeting mCherry described above, we confirmed that *hf*Cas13d indeed has diminished collateral activity (**Suppl. Figure 4a**). However, we also noted that *hf*Cas13d exhibited impaired on-target cleavage compared to ancestral *Rfx*Cas13d when using the same mCherry-targeting crRNA (**Figure 1d**). To more accurately compare the on-target and collateral activities of these two Cas13d variants, we titrated the concentrations of either the Cas13d effector or the crRNA in HEK293T cells, which led to dose-dependent silencing of mCherry (target) and EGFP (non-target). In addition to its compromised collateral activity, *hf*Cas13d showed a consistent reduction of its on-target cleavage activity across various concentrations of either the nuclease or crRNA tested (**Suppl. Figure 4b, 4c**). Our data therefore suggests that the mutagenesis of *hf*Cas13d, which eliminated its collateral activity, also impaired its on-target cleavage capability as compared with its predecessor, *Rfx*Cas13d. This observation differs from the findings reported in the original *hf*Cas13d study [34].

### *Rfx*Cas13d’s collateral activity exhibits negligible impact on endogenous RNA and protein homeostasis

Based on the results described above, we predicted that Cas13’s collateral activity would cleave proximal, endogenous RNAs, which may lead to global protein downregulation and loss of cell viability. To probe this hypothesis, we transfected HEK293T cells with plasmids encoding *Lwa*Cas13a, *Psp*Cas13b, or *Rfx*Cas13d orthologs together with either a non-targeting or mCherry-targeting crRNA and conducted mass spectrometry analysis for comparative proteome profiling between targeting and non-targeting conditions. In this proteomic analysis, *Lwa*Cas13a and *Psp*Cas13b nucleases exhibited potent silencing of the mCherry target without any notable degradation of GFP, *Lwa*Cas13a, *Psp*Cas13b, or other cellular proteins, confirming that these two Cas13 nucleases are highly potent and lack any collateral activity against exogenous or endogenous RNA in HEK293T cells (**Figure 1f, 1g**). *Rfx*Cas13d, however, showed a surprising proteome signature. First, proteomic analysis confirmed efficient silencing of the target mCherry and a significant collateral downregulation of the non-target GFP and *Rfx*Cas13d itself (**Figure 1h**), consistent with fluorescence and western blot data described above (**Figure 1d, 1e**). However, only a few endogenous cellular proteins exhibited modest downregulation due to *Rfx*Cas13d’s collateral activity (**Figure 1h)**. To better document the collateral activity of *Rfx*Cas13d, we conducted transcriptomic analysis. This analysis supported the proteomic findings, showing that the collateral activity primarily affects the RNA integrity of exogenous transcripts, with minimal impact on endogenous RNAs (**Figure 1i**).

We then investigated whether these findings could be replicated in other cell lines. To do this, we conducted similar experiments using HCT116 and HeLa cells. Results from fluorescence analysis, western blotting, and proteomics consistently showed that *Lwa*Cas13a and *Psp*Cas13b exhibited no collateral activity, whereas *Rfx*Cas13d’s collateral activity selectively degraded only overexpressed exogenous RNA (**Suppl. Figure 5**). This result indicated that the collateral activity of *Rfx*Cas13d selectively degrade ectopically overexpressed RNAs and has minimum impact on endogenous RNAs (**Suppl. Figure 5g**).

Since the collateral activity of *Rfx*Cas13d selectively alters exogenous RNA and proteins, we hypothesised that this would have minimal or no impact on cell fitness or survival. To test this, we measured cell metabolism and viability using a Resazurin assay. We assessed the viability of HEK293T cells expressing *Rfx*Cas13d in the presence of either mCherry-targeting and non-targeting crRNAs. As controls, we used *hf*Cas13d which has compromised on-target and collateral activities [34], and a catalytically inactive (dead) d*Rfx*Cas13d [42] (**Suppl. Figure 6a)**. The results showed similar cell metabolism across cells expressing *Rfx*Cas13d, dead d*Rfx*Cas13d, or *hf*Cas13d, regardless of the crRNA used. A flow-cytometry viability assay also confirmed that the *Rfx*Cas13d’s collateral activity does not cause any substantial cell death (**Suppl. Figure 6b**). Together, these findings indicate that, under these experimental conditions, the collateral activity of *Rfx*Cas13d does not impact cell metabolism or viability.

### The collateral activity of *Rfx*Cas13d is dependent on the expression level of the target

Next, we hypothesised that *Rfx*Cas13d’s on-target and collateral activity might be influenced by the intracellular abundance of *Rfx*Cas13d enzyme, its crRNA, the non-target transcript, or the target transcript, as these factors can dictate the abundance of *Rfx*Cas13d enzymes with activated nuclease domains. To test this hypothesis, we used the fluorescence-based assay described above to probe the collateral activity while systematically increasing the intracellular concentrations of *Rfx*Cas13d, crRNA, non-target (EGFP), or target (mCherry) transcripts via titration transfections of different plasmids. As expected, both on-target and collateral activities were highly dependent on the expression levels of *Rfx*Cas13d and its crRNA (**Suppl. Figure 7a**). In contrast, the intracellular abundance of the non-target EGFP had no effect on either on-target or collateral activity (**Suppl. Figure 7a**). Interestingly, only collateral degradation of EGFP was strongly correlated with the intracellular abundance of the target transcript mCherry, while on-target mCherry silencing remained potent regardless of target abundance (**Figure 2a**). High levels of target expression resulted in the most pronounced collateral degradation of EGFP (**Figure 2a, Suppl. Figure 7b-7d**). Western blot analysis further confirmed a positive correlation between target abundance and *Rfx*Cas13d self-cleavage through its collateral activity (**Figure 2b**). In contrast, altering the abundance of the non-target EGFP had no effect on the extent of collateral cleavage (**Suppl. Figure 8a-8e**).

**Figure 2.**
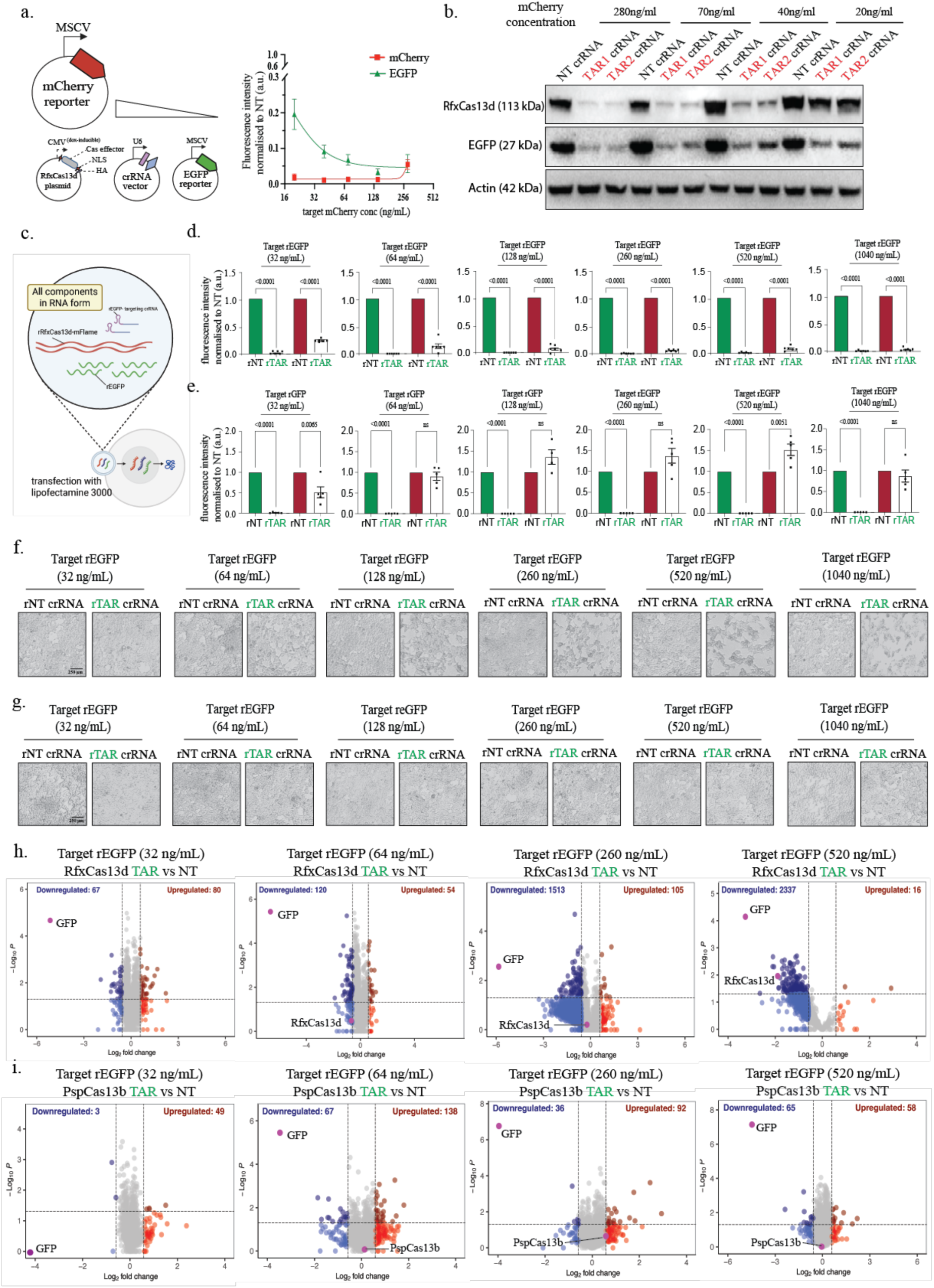
**a,** Quantification of the comparison of on-target (mCherry) and collateral cleavage (EGFP) silencing with a titration of mCherry plasmid. Data points in the graph are averages of the normalised mean fluorescence from four representative fields of view imaged (n = 3). **b,** Representative Western blot of the expression level of RfxCas13d, EGFP, and β actin protein levels in HEK293T cells expressing either mCherry-targeting crRNA or NT crRNA when titrating the transfected target mCherry plasmid. **c,** Schematics of RNA transfection of target rEGFP, rRfxCas13d-mFlame or rPspCas13b-mFlame, and rNT or rEGFP-targeting crRNA with lipofectamine 3000 into HEK293T cells. The ‘r’ prefix signifies the component was delivered in RNA form to distinguish from previous plasmid delivery forms. d, Quantification of RfxCas13d silencing of EGFP (target) or mFlame (non-target) transcripts with either rNT or rEGFP-targeting crRNAs at different target abundance (32ng/mL to 1040ng/mL). Data points in the graph are averages of the normalized mean fluorescence from four representative fields of view imaged (n = 6). The data are represented in arbitrary units (a.u.). Errors are the s.e.m. and p values of Student’s t-test are indicated (95% confidence interval). **e,** Quantification of PspCas13b silencing of EGFP (target) or mFlame (non-target) transcripts with either rNT or rEGFP-targeting crRNAs at different target abundance (32ng/mL to 1040ng/mL). Data points in the graph are averages of the normalized mean fluorescence from four representative fields of view imaged (n = 4 or 5). The data are represented in arbitrary units (a.u.). Errors are the s.e.m. and p values of Student’s t-test are indicated (95% confidence interval). **f&g,** Representative bright-field images of HEK293T cells 48 hours post-transfection of rRfxCas13d-mFlame or rPspCas13b-mFlame, rEGFP, and either rNT or rEGFP-targeting crRNA. (n= 5). Target EGFP RNA is transfected at a titration of different concentrations ranging from 32ng/mL to 1040ng/mL. **h&i,** Volcano plots showing the proteome of cells expressing EGFP, EGFP-targeting or NT crRNA paired with RfxCas13d-mFlame or PspCas13b-mFlame in HEK293T cells at four different target EGFP concentration (32, 64, 260, 520 ng/mL). Data points represent proteins, with log_2_FC > 0.58 (upregulation, blue) or log_2_FC < −0.58 (downregulation, red) and p < 0.05 indicating significantly differential expression. Total downregulated and upregulated proteins are shown at the top corners. Exogenous Cas13 and EGFP are labelled purple. Results were analysed by an unpaired two-tailed Student’s t-test (n = 5).

Together, these findings underscore the pivotal role of target transcript abundance in amplifying *Rfx*Cas13d activation, thereby driving increased collateral activity in human cells.

### Elevated target RNA levels drive global deregulation of endogenous proteins via amplified collateral activity

To further investigate the impact of target abundance on collateral activity, we shifted from a plasmid to RNA delivery. We reasoned that, unlike plasmids, RNA transfection would enable immediate translation of *Rfx*Cas13d and rapid availability of the target RNA, offering better temporal control to study the dynamics of collateral activity. Indeed, RNA transfection into HEK293T cells produced over 250 times more target RNA than plasmid transfection within just six hours, confirming that direct RNA delivery can offer a rapid availability of target RNA with high abundance (**Suppl. Figure 9**).

We first generated an *Rfx*Cas13d-mFlame mRNA, where the mFlame fluorescence serves as a surrogate marker for both RfxCas13d expression and its collateral activity upon transfection into HEK293T cells (**Figure 2c**). To assess whether target abundance impacts collateral activity of *Rfx*Cas13d, we co-transfected HEK293T cells with increasing concentrations of EGFP target mRNA along with constant concentration of *Rfx*Cas13d-mFlame mRNA and crRNAs. We monitored EGFP (target) and mFlame (non-target) fluorescence, along with cell morphology, at 4-, 24- and 48-hours post-transfection (**Suppl. Figure 10**).

As expected, the EGFP-targeting crRNA caused a strong repression of both EGFP and mFlame fluorescence across all tested target RNA concentrations compared to the non-targeting crRNA control, validating the functionality of *Rfx*Cas13d in this RNA delivery setting (**Figure 2d, Suppl. Figure 10**). Notably, we observed significant changes in cell morphology when *Rfx*Cas13d was programmed to silence highly abundant EGFP target RNA (**Figure 2f, Suppl. Figure 10b, 10c**). Brightfield images revealed cellular toxicity, suggesting that high target abundance leads to excessive *Rfx*Cas13d activation and subsequent degradation of host RNA as a result of heightened collateral activity. Resazurin and CellTiter-Glo assays confirmed substantial impairment of cell metabolism and proliferation when the nuclease domains of *Rfx*Cas13d are activated by high levels of target RNA, whereas low target abundance had no or very limited effect on cell metabolism and proliferation (**Suppl. Figure 11a, 11b**). In contrast, Zombie NIR assay exhibited no effect of RfxCas13d and target abundance on cell death (**Suppl. Figure 11c-11f**). We obtained similar results when RfxCas13d was delivered through lentiviral transduction while crRNA and target RNA were delivered by RNA transfection (**Suppl. Figure 12**). This data suggests that the collateral activity triggers cell dormancy rather than cell death in HEK293T cells.

To further characterize the impact of collateral activity on cellular homeostasis, we performed mass spectrometry-based proteomics to analyse changes in protein expression under varying conditions of target RNA abundance. Our analyses revealed that increasing target RNA abundance correlates with broader alterations in the proteome, consistent with heightened *Rfx*Cas13d activation and collateral degradation of host RNA (**Figure 2h**). Notably, these proteomic changes were more pronounced under conditions of high target RNA abundance, aligning with our observations of impaired cell morphology, metabolism, and proliferation. In contrast, cells transfected with low levels of target RNA exhibited minimal changes in protein expression, suggesting limited collateral activity. We performed similar experiments using *Psp*Cas13b-mFlame mRNA and observed minimal changes in cell morphology and proteome expression across different target concentrations (**Figure 2e, 2g, 2i, Suppl. Figure 13**). This data aligns with the lack of collateral activity associated with *Psp*Cas13b described above.

### Endogenous transcripts with moderate expression restrict the collateral activity of *Rfx*Cas13d

Previously, we assessed Cas13 collateral activity using exogenous targets ectopically expressed in human cells. However, ectopic expression of exogenous RNA is known to differ significantly from endogenous transcripts in terms of expression levels and cellular localisation [43–45], which could influence the collateral activity. We investigated whether targeting endogenous transcripts could modulate the collateral activity of different Cas13 orthologs. We used *Lwa*Cas13a, *Psp*Cas13b, *Rfx*Cas13d, and their respective crRNAs encoded in plasmids to target the transcript of beta-2-microglobulin (β2m) in HEK293T cells (**Figure 3a**). As a non-essential protein within the MHC class I complex, β2m was selected because its silencing is predicted to have negligible impact on broader gene and protein expression beyond the MHC class I system. To monitor the extent of collateral activity when targeting β2m transcript, we used EGFP and BFP fluorescence encoded in the Cas13 plasmid constructs as surrogate markers. FACS analysis confirmed efficient silencing of β2m by all three Cas13 orthologs (**Figure 3b**). Consistent with our previous observations, *Lwa*Cas13a and *Psp*Cas13b did not exhibit any collateral activity, however, *Rfx*Cas13d displayed only a moderate collateral activity against the non-target RNA reporters, which was not statistically significant (**Figure 3b)**.

**Figure 3.**
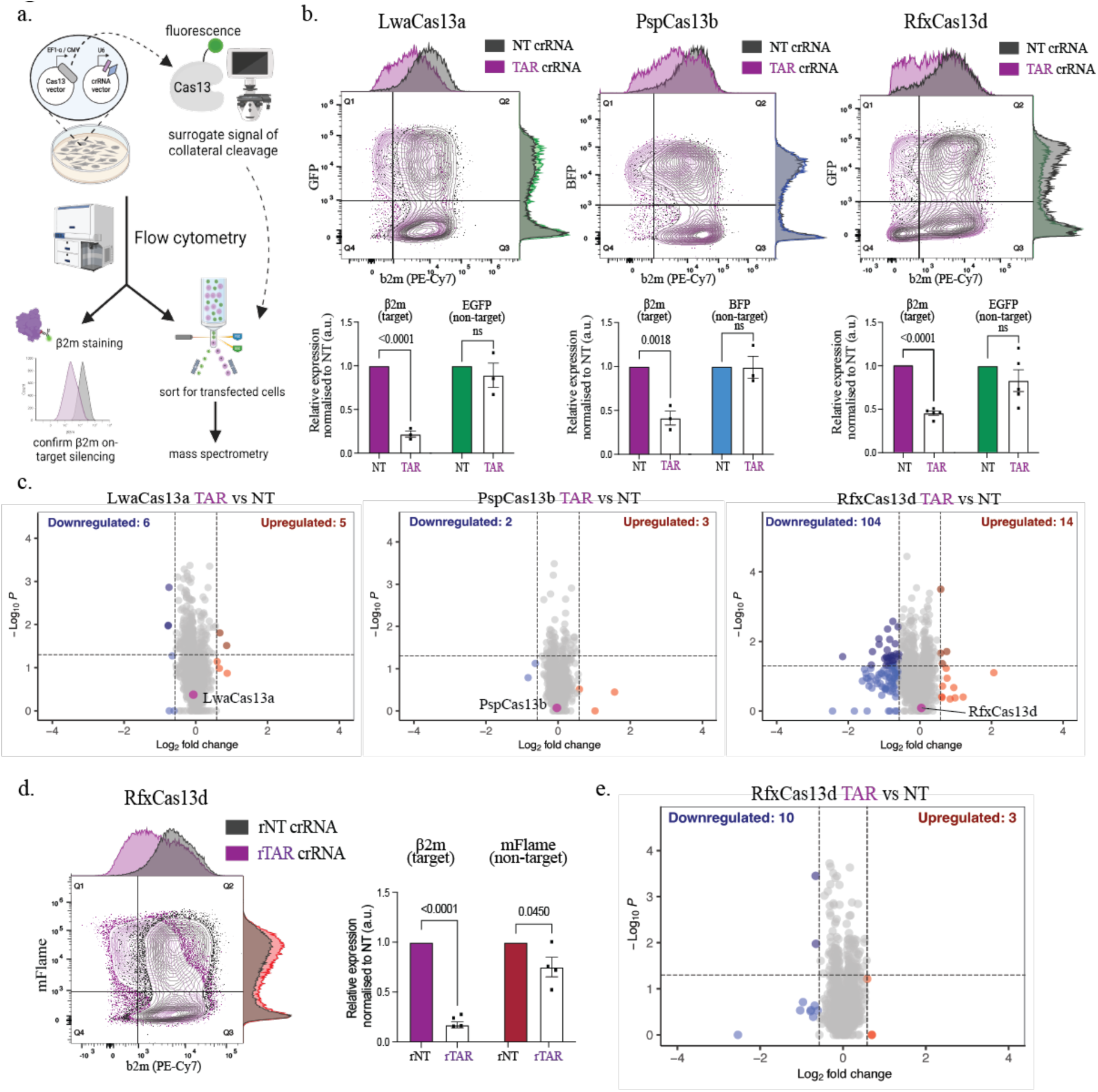
**a,** Schematic of FACS sorting of Cas13 positive cells after Cas13 and crRNA plasmid transfection endogenous β2m silencing assay, and subsequent protein extraction for downstream proteomic analyses. **b, Top,** Representative FACS contour plots showing the level of β2m and fluorescent protein tagged on Cas13 effectors expression at 72 hours post transfection in HEK 293T cells. The histogram on the x-axis shows the silencing of endogenous β2m expression (on-target cleavage) for β2m-targeting crRNA (purple) compared to NT crRNA (black). The histogram on the y-axis shows the silencing of the fluorescent EGFP protein expression (collateral cleavage) for β2m-targeting crRNA compared to NT crRNA (n=3). **Bottom,** Quantification of Cas13 silencing of β2m (on-target cleavage) and fluorescence proteins (collateral cleavage) with β2m-targeting crRNA compared to NT crRNA. Errors are the s.e.m. and p values of Student’s t-test are indicated (95% confidence interval). **c,** Volcano plots showing the proteome of cells expressing β2m-targeting or NT crRNA paired with different Cas13 orthologs in HEK293T cells. Data points represent proteins, with log_2_FC > 0.58 (upregulation, blue) or log_2_FC < −0.58 (downregulation, red) and p < 0.05 indicating significantly differential expression. Total downregulated and upregulated proteins are shown at the top corners. Exogenous Cas13 is labelled purple. Results were analysed by an unpaired two-tailed Student’s t-test (n = 5). **d**, **Left**, Representative FACS contour plots showing the level of β2m and fluorescent protein tagged on Cas13 effectors expression at 48 hours post transfection in HEK293T cells. The histogram on the x-axis shows the silencing of endogenous β2m expression (target) for rβ2m-targeting crRNA (purple) compared to non-targeting (rNT) crRNA (black). The ‘r’ prefix signifies the component was delivered in RNA form. The histogram on the y-axis shows the silencing of the fluorescent mFlame protein expression (collateral cleavage) for β2m-targeting crRNA compared to NT crRNA (n=5). **Right**, Quantification of Cas13 silencing of β2m (target) and fluorescence proteins (collateral cleavage) with β2m-targeting crRNA compared to NT crRNA. Errors are the s.e.m. and p values of Student’s t-test are indicated (95% confidence interval). **e,** Volcano plots showing the proteome of cells expressing rβ2m-targeting or rNT crRNA paired with different Cas13 orthologs in HEK293T cells. Data points represent proteins, with log_2_FC > 0.58 (upregulation, blue) or log_2_FC < −0.58 (downregulation, red) and P < 0.05 indicating significantly differential expression. Total downregulated and upregulated proteins are shown at the top corners. Cas13 is labelled purple. Results were analysed by an unpaired two-tailed Student’s t-test (n = 5).

We conducted quantitative proteomic analyses to assess the impact of collateral activity on endogenous protein expression during β2m targeting. Although mass spectrometry-based quantitative proteomics could not detect the target β2m due to its moderate expression in HEK293T cells, its silencing was validated by FACS analysis, confirming the activation of HEPN domains that is essential for the collateral activity to occur (**Figure 3b**). LwaCas13a and PspCas13b did not show any alteration in protein expression due to the collateral activity, consistent with our prior findings. In contrast, RfxCas13d’s collateral activity exhibited only a moderate downregulation of a limited number of endogenous proteins, and no self-silencing of its own RfxCas13d protein (**Figure 3c**). We obtained similar results where no collateral cleavage at a global cellular level was detected when RfxCas13d-mFlame and its crRNAs were delivered in RNA form (**Figure 3d, 3e**).

We assessed whether these characteristics of collateral activity persist when targeting endogenous β2m transcript across human cell lines using lentiviral delivery of RfxCas13d and its crRNA. First, HEK293T cells were sequentially transduced with doxycycline (dox)-inducible RfxCas13d and crRNA lentiviruses, and collateral activity was assessed upon targeting endogenous β2m mRNA. FACS analysis showed efficient, dox concentration–dependent inhibition of β2m protein expression (**Suppl. Figure 14a**). Consistent with our previous findings, mass spectrometry–based proteomics revealed no evidence of collateral activity despite robust target silencing across dox concentrations (**Suppl. Figure 14b**). Similar proteomic analyses in RPMI and OPM2 cell lines, which express levels of β2m protein detectable by mass spectrometry, confirmed potent silencing of β2m and associated HLA subunits of the MHC class I complex, while again showing no global protein deregulation, supporting no or minimal collateral activity of RfxCas13d when targeting this endogenous transcript (**Suppl. Figure 14c, 14d**).

Together, these findings underscore that intracellular RNA abundance is a critical determinant of RfxCas13d collateral activity in human cells. While moderately abundant endogenous targets induce no or limited collateral effects, highly abundant exogenous targets activate a greater proportion of RfxCas13d molecules, which compromise intracellular RNA and protein homeostasis.

### Target abundance controls collateral activity in zebrafish embryos

The data above demonstrate that target RNA abundance dictates the extent of collateral cleavage by RfxCas13d. We next asked whether these collateral cleavage properties observed in human cell lines also hold true in vivo. To investigate this, we utilized the zebrafish model, where RfxCas13d has been previously shown to effectively knock down gene expression [41, 46]. We reasoned that on-target cleavage of exogenous EGFP transcript by RfxCas13d would not affect embryonic development, as EGFP is dispensable. However, extensive collateral cleavage of maternal and zygotic RNAs would be expected to impair development. This model further allows us to titrate target abundance through controlled RNA injection and directly assess developmental defects arising from collateral activity. We injected RfxCas13d-mFlame, rEGFP, and crRNAs non-targeting (rNT crRNA) or targeting EGFP (rEGFP-targeting crRNA) in RNA form into one-cell-stage zebrafish embryos and used the same dual-fluorescence system to assess on-target and collateral cleavage (**Figure 4a**). We used fluorescence microscopy to monitor the expression of EGFP (on-target) and RfxCas13d-mFlame (collateral) in vivo at 4 hours post-fertilization (hpf) and 1 day post-fertilization (dpf).

**Figure 4.**
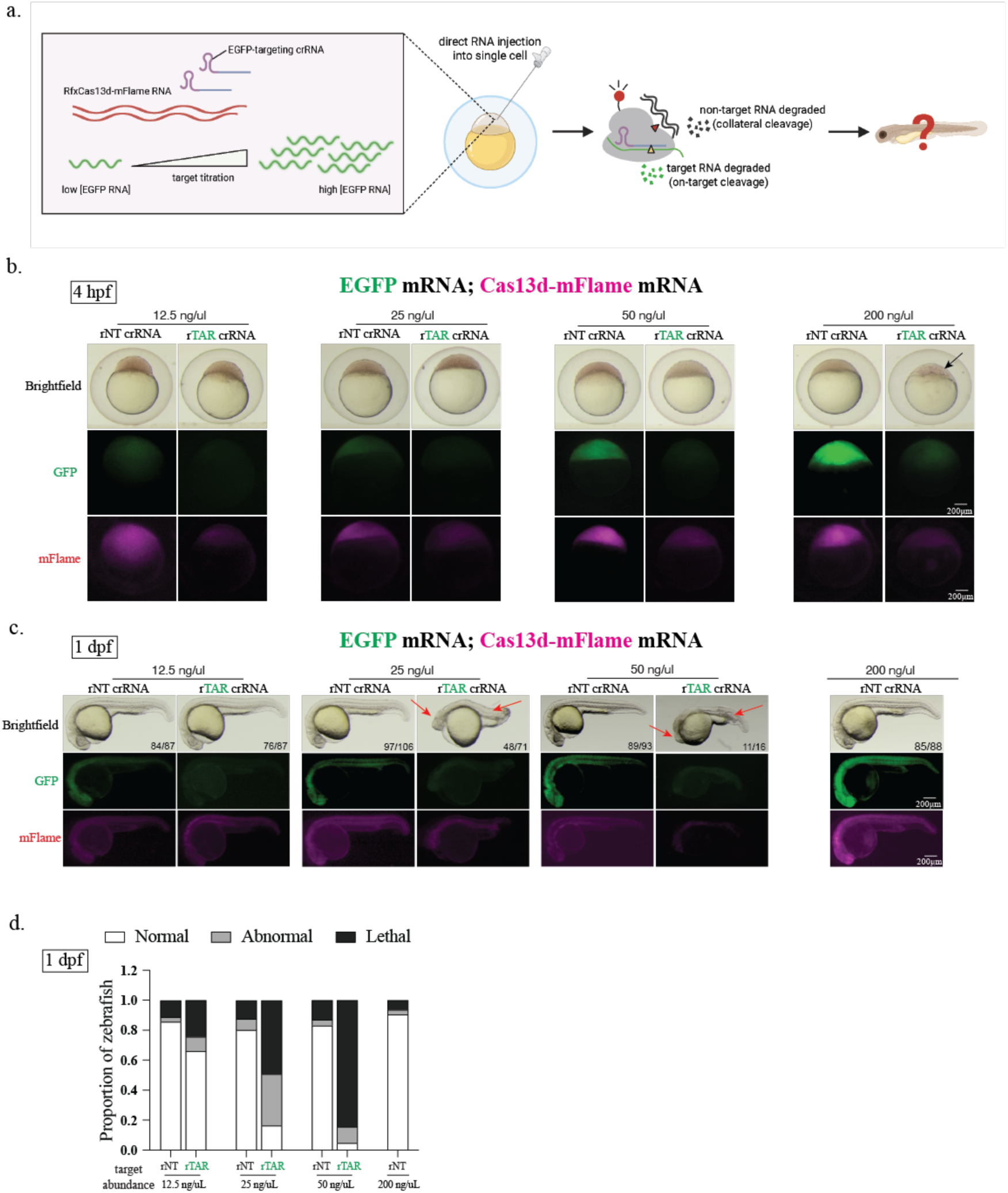
**a**, Schematic of RNA injection of rRfxCas13d-mFlame, rEGFP, and rEGFP-targeting or non-targeting (rNT) control crRNA into zebrafish embryo at different titration of target EGFP RNA concentration. **b**, Bright-field and fluorescent images of representative embryos at 4 hpf (hours post-fertilisation) showing embryo morphology as well as EGFP on-target cleavage (green) and mFlame collateral cleavage (magenta) when comparing rEGFP-targeting crRNA to rNT crRNA at four different target rEGFP concentrations (12.5, 25, 50 and 200 ng/μL). The arrow indicates developmental abnormality. Scale bar is 200μm. **c,** Bright-field and fluorescent images of representative embryos at 1 dpf (day post-fertilisation) showing embryo morphology as well as EGFP on-target cleavage (green) and mFlame collateral cleavage (magenta) when comparing EGFP-targeting crRNA to NT crRNA at four different target rEGFP concentrations (12.5, 25, 50 and 200 ng/μL). Numbers on the bottom right corner shows representative zebrafish out of all fertilised and survived zebrafish embryos. Scale bar is 200μm. **d,** Quantification of the proportion of zebrafish that are normal (white), abnormal (grey) and lethal (black) at 1 dpf after injection with rEGFP mRNA at four different concentrations (12.5, 25, 50 and 200 ng/μL).

At 4 hpf, embryos injected with rEGFP-targeting crRNA exhibited strong silencing of both EGFP (target) and mFlame (non-target), whereas the EGFP and mFlame fluorescence in embryos injected with the non-targeting crRNA remained high (**Figure 4b**). This result suggests that the RfxCas13d HEPN domains were activated in the embryos and mediated the degradation of the target EGFP RNA, whereas the collateral activity impaired mFlame expression. However, while the collateral cleavage is activated at all concentrations, we noticed that, only when target EGFP RNA was injected at the highest quantity (200 pg, 200ng/μL), 95.6% of zebrafish embryos exhibited obvious developmental defects at 4 hpf and none survived to 1 dpf, **(Figure 4b**). When the target EGFP RNA decreases in quantity, the percentage of abnormal or dead zebrafish at 1 dpf decreased accordingly. For instance, the majority (83.7%) of zebrafish embryos injected with the lowest target EGFP RNA concentration (12.5 ng/μL) exhibited only mild pericardial oedema formation at 1 dpf (**Figure 4c, 4d**). All tested RNA target concentrations appeared to be embryonic lethal beyond 1 dpf, but the severity of developmental defects is proportional to target RNA concentrations, consistent with earlier lethality at the highest concentration. These results further validated that collateral cleavage activities are positively correlated with target RNA abundance.

### Target localisation drives tissue-specific collateral activity and developmental defects

We then tested if the collateral activity of RfxCas13d would affect zebrafish development when the target EGFP is expressed under tissue-specific promoters at later developmental stages, even when RfxCas13d and its crRNA are injected at single-cell stage. We hypothesised that collateral activity-induced developmental defects would be confined to the specific tissues expressing the target rEGFP. To test this, we used the Tg(fli1a:nEGFP) transgenic zebrafish, which endogenously express EGFP in endothelial progenitors starting at ∼12 hours post-fertilisation (hpf) [47, 48]. Heterozygous Tg(fli1a:nEGFP) transgenic zebrafish was crossed with wildtype (AB) to generate siblings with 50% Tg(fli1a:nEGFP) transgenic and 50% AB. We observed that ∼95% of Tg(fli1a:nEGFP) transgenic embryos developed normally at both 1 and 2 dpf in rNT crRNA condition. Similarly, around 93% of AB embryos (without target EGFP) developed normally at both 1 and 2 dpf in the rEGFP-targeting condition. In comparison, we observed significant downregulation of endothelial EGFP expression in Tg(fli1a:nEGFP) transgenic embryos in EGFP-targeting condition at both 1 and 2 dpf, and also noted 51% of embryos displayed significant pericardial oedema at 2 dpf (**Figure 5a**). This shows that only embryos endogenously expressing the target EGFP exhibited tissue-specific developmental defects in EGFP-targeting condition. To confirm that this result is not limited to a single endothelial promoter, we also tested this in embryos generated from crossing of heterozygous Tg(kdrl:EGFP) transgenic zebrafish with AB adults. Like the fli1a promoter, the kdrl promoter drives EGFP expression from ∼13 hpf in endothelial progenitors [48]. All EGFP-positive embryos developed normally under the rNT crRNA condition, and nearly all AB embryos lacking EGFP developed normally under the rEGFP-targeting crRNA condition at 1 and 2 dpf. In contrast, ∼90% of EGFP-expressing embryos displayed significant pericardial oedema at 2 dpf under the rEGFP-targeting crRNA condition (**Figure 5b**). This pericardial oedema phenotype was accompanied by robust silencing of kdrl:EGFP expression at both 1 and 2 dpf. Collateral activity in both Tg(fli1a:EGFP) and Tg(kdrl:EGFP) transgenics resulted in pericardial oedema indicative of cardiovascular defects, with mild overall phenotype when compared to EGFP mRNA-injected embryos (**Figure 4**). This shows that localized expression of abundant target RNA leads to tissue-specific collateral activity and moderate developmental abnormalities (**Figure 5a, 5b**).

**Figure 5.**
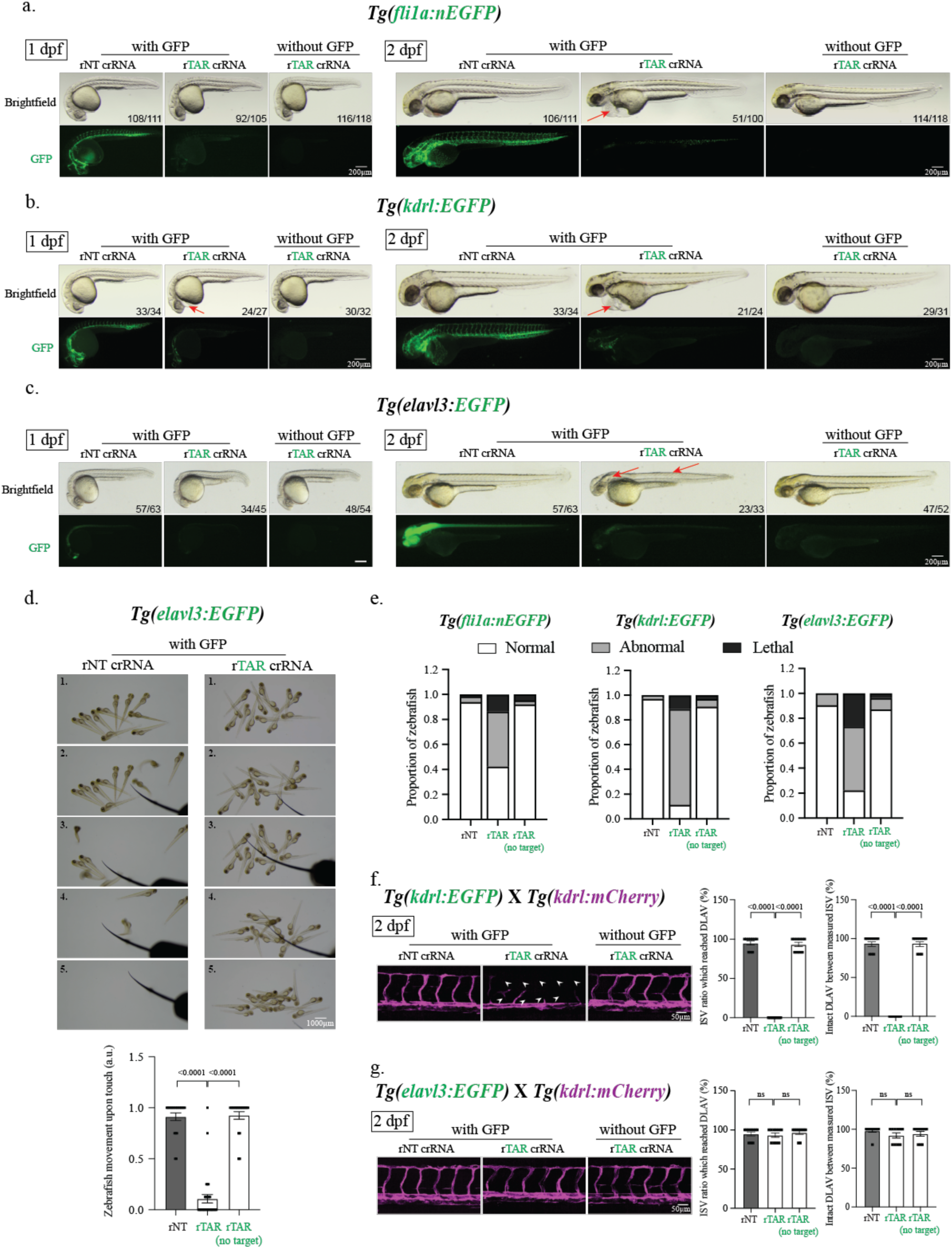
**a-c,** Brightfield and fluorescent images of **a,** Tg(fli1a:nEGFP), **b,** Tg(kdrl:EGFP), and **c,** Tg(elavl3:EGFP) transgenic embryos showing embryo morphology as well as EGFP on-target cleavage using RfxCas13d with EGFP-targeting crRNA or NT crRNA control at 1 and 2 dpf (days post-fertilisation). Sibling embryos from the same spawning batch without EGFP expression were also injected with RfxCas13d with EGFP-targeting crRNA as a negative control. Arrows indicate developmental abnormalities (pericardial oedema in **a, b** brain and neural tube area deformities in **c**). Numbers on the bottom right corner shows representative zebrafish out of all fertilised and survived zebrafish embryos. Scale bar is 200μm. **d, Top**, Representative snapshot images showing movement of 3 dpf Tg(elavl3:EGFP) zebrafish larvae injected with rRfxCas13d-mFlame paired with either rNT or rEGFP-targeting crRNA. Scale bar is 1mm. **Bottom**, Quantification of larvae movement (a.u.) upon touch in Tg(elavl3:EGFP) or wild type (AB) sibling larvae injected with rRfxCas13d-mFlame paired with either rNT or rEGFP-targeting crRNA. Scale bar is 1000μm. **e,** Quantification of the proportion of zebrafish that are normal (white), abnormal (grey) and lethal (black) after injection at 2dpf for different transgenic lines. **f-g, Left,** Lateral spinning disc confocal images (magenta showing kdrl:mCherry expression) of **f,** Tg(kdrl:EGFP); Tg(kdrl:mCherry) or **g,** Tg(elavl3:EGFP); Tg(kdrl:mCherry) double transgenic embryos at 2dpf. Lack of intersegmental vessel (ISV) and dorsal longitudinal anastomotic vessel (DLAV) development indicated with white arrow heads. Scale bar is 50μm. **Right,** Quantification of complete ISV (ratio of ISV that have reached the DLAV level) and DLAV (ratio of fully connected DLAV between ISVs) formation.

Interestingly, comparison of EGFP transcript levels at 1 dpf revealed that wildtype embryos injected with 200 pg of exogenous EGFP mRNA exhibited the highest EGFP abundance and the most severe developmental defects, whereas Tg(fli1a:nEGFP) and Tg(kdrl:EGFP) transgenic embryos showed lower transcript levels with correspondingly milder abnormalities. These results suggest that target abundance, localisation, and timing of expression determine the severity of collateral cleavage–induced developmental defects (**Suppl. Figure 15a,** **Figure 4b, 5a, 5b**)

### RfxCas13d collateral activity in neurons induces CNS abnormalities and mobility deficits

We next investigated whether EGFP expression in neurons could trigger localized developmental defects in the central nervous system and impair zebrafish mobility without causing pericardial oedema. To test this, we crossed heterozygous Tg(elavl3:EGFP) transgenic zebrafish, in which the elavl3 promoter drives neuronal EGFP expression in the brain and neural tube from ∼10 hpf [49]. Approximately 91–93% of embryos developed normally under control conditions, including EGFP-expressing zebrafish injected with rNT crRNA and EGFP-negative zebrafish injected with rEGFP-targeting crRNA at 1 and 2 dpf. In contrast, elavl3:EGFP-positive embryos injected with rEGFP-targeting crRNA displayed pronounced developmental defects in the brain and neural tube region at 2 dpf, accompanied by marked silencing of elavl3:EGFP expression at both 1 and 2 dpf (**Figure 5c)**. At 3 dpf, Tg(elavl3:EGFP) transgenic larvae injected with RfxCas13d and rEGFP-targeting crRNA exhibited lethargy and impaired movement, phenotypes observed exclusively in EGFP-positive fish and consistent with the neurological defects identified in these zebrafish (**Figure 5d**, **Suppl. Video 1-3**). Importantly, unlike vascular EGFP reporter transgenics (i.e., fli1a:nEGFP and kdrl:EGFP) (**Figure 5a, 5b**), collateral cleavage in elavl3:EGFP transgenics did not result in pericardial oedema (**Figure 5c**). When quantifying and comparing all three different crosses of Tg(fli1a:nEGFP), Tg(kdrl:EGFP) and Tg(elavl3:EGFP) transgenics at 2dpf, we noticed a similar trend where the EGFP-targeting crRNA activated RfxCas13d to cause collateral cleavage that led to substantial developmental defect or death, which did not occur for rNT crRNA or for when target EGFP was absent (**Figure 5e**).

To further understand the difference in morphological abnormalities caused by RfxCas13d-induced collateral cleavage between neuronal or endothelial EGFP reporter transgenics, we compared their vascular development using confocal microscopy. When closely examining the intersegmental vessel (ISV) and the dorsal longitudinal anastomotic vessel (DLAV) development between 2 dpf Tg(elavl3:EGFP) and Tg(kdrl:EGFP) transgenic embryos injected with either rNT crRNA or rEGFP-targeting crRNA, we observed significant impairment of ISV and DLAV development only in Tg(kdrl:EGFP) transgenic embryos under the rEGFP-targeting condition (**Figure 5f**, **Suppl. Figure 15b**). In comparison, we observed relatively normal ISV and DLAV development in Tg(elavl3:EGFP) transgenic embryos under the rEGFP-targeting condition, except for the reduced ISV length likely due to neural tube abnormalities causing reduced trunk size (**Figure 5f, 5g, Suppl. Figure 15c**).

These data together show that RfxCas13d collateral cleavage is highly localised in zebrafish, and the target expression site dictates the location where collateral cleavage exerts its effects, leading to varying developmental abnormalities.

## DISCUSSION

Our study identifies key fundamental mechanisms that govern on-target and collateral RNA cleavage activities of diverse Cas13 orthologs. We demonstrated that LwaCas13a and PspCas13b orthologs are highly specific and lack detectable collateral RNA degradation across multiple human cell lines. Their lack of collateral activity makes them ideal programmable nucleases for RNA silencing applications, including forward genetic screens of non-coding RNA [50, 51], which could further improve the accuracy and quality of these screens. Unexpectedly, we found that the engineered high-fidelity Cas13d (hfCas13d) exhibited an impaired on-target activity likely due to the mutagenesis that inactivated its collateral activity. These mutations yielded marked reduction in RNA silencing compared to its ancestor RfxCas13d. This observation of attenuated on-target activity of hfCas13d aligns with findings from a recent study [51], but contrasts with the initial report, which described hfCas13d as retaining unaltered on-target activity [34].

Among the Cas13 orthologs we investigated, RfxCas13d exhibited a collateral activity with surprising properties. We found that when targeting RNA transcripts with low to moderate intracellular abundance, RfxCas13d exhibits a collateral activity that predominantly degraded ectopically expressed transcripts but had no or very little impact on endogenous protein homeostasis or cell fitness. However, RNA targets with high abundance unleashed marked collateral activity that altered proteomic homeostasis and led to significant cell toxicity. We propose that the low to moderate target abundance is likely to activate the HEPN nuclease domains of only a small fraction of RfxCas13d molecules that are available in the cell, leaving the vast majority inactive (**Figure 6**). The few target-activated RfxCas13d enzymes have a low probability of encountering other endogenous cellular RNAs, limiting the scale of collateral RNA degradation and cell toxicity. Conversely, when a target is highly abundant, a simultaneous activation of high number of RfxCas13d molecules would take place, which significantly heightens the probability of encounter with essential cellular RNA and mediate their degradation as collateral effect of HEPN nuclease domains activation (**Figure 6**). Our data suggest that target RNA abundance needs to reach high levels to elicit detectable effects of collateral cleavage on cell fitness. This implies that when utilising RfxCas13d to silence endogenous transcripts with moderate expression levels, target RNA abundance is likely to be a limiting factor to prevent collateral cleavage from occurring at a significant level. This is supported by the β2m-silencing proteomics results, which showed no or limited collateral activity despite efficient silencing of β2m target mRNA.

**Figure 6.**
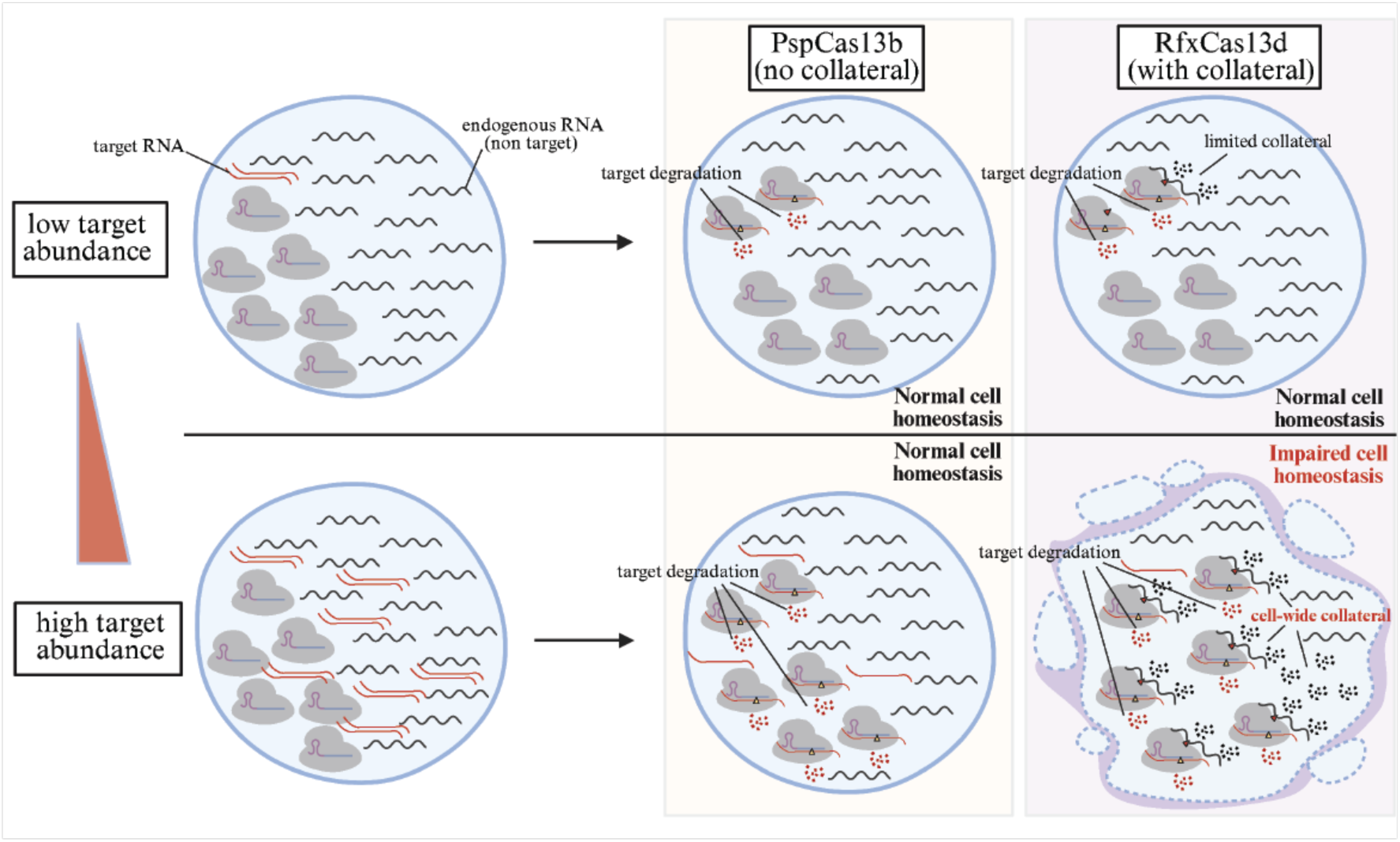
Schematics of the proposed model illustrating how target RNA abundance influences collateral RNA cleavage.

A surprising finding of this study is the heightened sensitivity of ectopically expressed RNAs to RfxCas13d collateral activity compared to endogenous transcripts. Notably, under moderate target expression, we observed robust degradation of exogenous RNA (e.g. reporter and RfxCas13d mRNA) mediated by the collateral activity of RfxCas13d, while endogenous RNAs and proteins persisted at steady levels, consistent with the absence of cellular toxicity. This selective degradation suggests that ectopically expressed RNAs may preferentially engage with activated RfxCas13d HEPN domains, potentially due to spatial proximity or elevated local concentrations. However, endogenous RNAs are less prone to degradation by RfxCas13d collateral activity, likely owing to protective cellular context or limited interaction with the nuclease domains. These findings align with previous reports showing that the collateral activity of certain Cas13 orthologs can induce a state of cellular dormancy in bacteria, driven by widespread non-specific cleavage of host RNAs, a process that is strongly dependent on the level of bacteriophage-derived transcripts in the infected cell [24, 52].

Our zebrafish data also indicate that the extent of RfxCas13d collateral activity depends on target RNA abundance. When the target RNA was expressed at low or moderate levels, zebrafish development was either unaffected or only mildly impacted. This is in accordance with a recent study where RfxCas13d was successfully used to silence maternal RNA in the embryo of zebrafish without significant collateral activity [46]. In contrast, the activation of RfxCas13d collateral activity in zebrafish embryos by a highly abundant target RNA caused significant developmental defect and animal death. The severe phenotype observed in the zebrafish is likely due to the collateral degradation of maternal and zygotic RNA molecules that are critical in coordinating normal embryonic development [53–56].

When we used transgenic zebrafish lines in which the expression of EGFP mRNA target is restricted to endothelial cells at a later developmental stage, we observed localised developmental abnormalities affecting the ISV and the DLAV, leading to pericardial oedema formation. Similarly, when target EGFP RNA expression is restricted to neurons, abnormal development occurred in the CNS region and impaired mobility of the zebrafish, without causing pericardial oedema. These data indicate that restricted expression of the target RNA in specific tissues at a later developmental stage elicit only localized developmental abnormalities. In contrast, target expression at single-cell stage causes a more pronounced and global developmental defects.

Together, our findings demonstrate that RfxCas13d exhibits potent collateral activity in mammalian cells and zebrafish embryos in a manner strongly dependent on target abundance, helping to resolve a longstanding controversy regarding the determinants of Cas13 collateral activity in eukaryotic cells and organisms [2, 29–37]. This target abundance-dependent collateral activity of RfxCas13d may be leveraged for selective elimination of virally infected or cancerous cells that express high levels of viral or aberrant RNA species. In contrast, PspCas13b and LwaCas13a display markedly reduced collateral activity, making them better suited for precise RNA knockdown applications.

## METHODS

### Design and cloning of crRNAs for LwaCas13a, PspCas13b, RfxCas13d

The design and cloning of PspCas13b, RfxCas13d mCherry-targeting crRNAs were according to a previous publication [13]. The LwaCas13a mCherry-targeting crRNAs were optimised via screening random spacer designs. Briefly, individual guide RNAs were cloned into the pC0043-PspCas13b crRNA backbone (addgene#103854, a gift from Feng Zhang lab, later referred to as crRNA backbone) which contains a PspCas13b crRNA direct repeat (DR) sequence and two BbsI restriction sites for the cloning of spacer sequence. A total of 20 μg crRNA backbone was digested by BbsI restriction enzymes (NEB, R3539) following the manufacturer’s instructions for 2 hours at 37°C. Backbone linearization was checked with 1% agarose gel. The digested backbone was purified with NucleoSpin Gel and PCR Clean-up Kit (Macherey-Nagel, 740609.50), aliquoted, and stored in −20°C until using.

For crRNA cloning, forward and reverse single-stranded DNA oligonucleotides containing CACC and CAAC overhangs respectively, were ordered from Sigma or IDT (100 μM). A total of 1.5 μL of 100 μM the forward and reverse DNA oligos were annealed in 47 μL annealing buffer (5 μl NEB buffer 3.1 and 42 μL H_2_O) by 5 min incubation at 95 °C and slow cool down in the heating block overnight. 1 μL of the annealed oligos were ligated with 0.04 ng digested PspCas13b crRNA backbone in 10 μL of T4 ligation buffer (3 h, RT) (Promega, M1801). All PspCas13b crRNA spacer sequences and target sequences used in this study are listed in **Supplementary Table 1**. All crRNAs and PspCas13b clones that are generated in this study were verified by Sanger sequencing (AGRF, AUSTRALIA). The primers used for PCR and Sanger sequencing are listed in Supplementary Table 3.

### Plasmid amplification and purification

The plasmid amplification and purification are described in a previous publication^18^. Briefly, TOP10 or Stbl3 bacteria were used for transformation. A total of 5–10 μL ligated plasmids were transformed into 30 μL of chemically competent bacteria by heat shock at 42 °C for 45 s, followed by 2 min on ice. The transformed bacteria were incubated in 500 μL LB broth media containing 75 μg/mL ampicillin (Sigma-Aldrich, A9393) for 1 h at 37 °C in a shaking incubator (200 rpm). The bacteria were pelleted by centrifugation at 6,000 rpm for 1 min at room temperature (RT), re-suspended in 100 μL of LB broth, and plated onto a pre-warmed 10 cm LB agar plate containing 75 μg/mL ampicillin, and incubated at 37 °C overnight. The next day, single colonies were picked and transferred into bacterial starter cultures and incubated for ∼6 h for mini-prep (Macherey-Nagel, NucleoSpin Plasmid Mini kit for plasmid DNA, 740588.50) or maxi-prep (Macherey-Nagel, NucleoBond Xtra Maxi Plus, 740416.50) DNA purification according to the standard manufacturer’s protocol.

### Cell culture

HEK293T (ATCC CRL-3216), HeLa, and HCT116 cell lines were cultured in DMEM high glucose media (Thermo Fisher, 11965092) containing 10% heat-inactivated fetal bovine serum (Thermo Fisher, 10100147), 100 mg/mL Penicillin/-Streptomycin (Thermo Fisher, 151401220), and 2 mM GlutaMAX (Thermo Fisher, A1286001). The cells were maintained at confluency between 20 and 80% in 37 °C incubators with 10% CO_2_.

RPMI and OPM2 cell lines were cultured in RPMI 1640 media (Thermo Fisher, 11875093) containing 10% heat-inactivated fetal bovine serum (Thermo Fisher, 10100147), 100 mg/mL Penicillin/-Streptomycin (Thermo Fisher, 151401220), and 2 mM GlutaMAX (Thermo Fisher, A1286001). The cells were maintained at confluency between 20 and 80% in 37 °C incubators with 10% CO_2_.

### RNA silencing assays using transfections

All DNA and most RNA transfection experiments were performed using an optimised Lipofectamine 3000 transfection protocol (Thermo Fisher, L3000015). For RNA silencing in HEK293T, cells were plated at approximately 120,000 cells per well (500uL) in 24-well plates (Corning) a day prior to transfection. Cells were transfected when they reached 70-90% confluency. For each well, a total of around 500 ng DNA plasmids or around 1000ng RNA were mixed with 1 μL P3000 reagent and 1.5 uL Lipofectamine 3000 in Opti-MEM Serum-free Medium (Thermo Fisher, 31985070) to a total of 53 uL. Each plasmid’s quantity used for each Cas13 orthologue’s silencing assay were previously optimised and listed below. Two targeting crRNA with the highest on-target cleavage were chosen and compared to one non-targeting crRNA as the negative control to establish relative silencing efficiency. The plasmid mixture were incubated for 20 min at room temperature and then added to each well for efficient delivery of plasmids. Transfected cells were incubated at 37°C and 10% CO2, and the transfection efficacy was monitored 24 and 48 hours post-transfection by fluorescent microscopy. Cells were harvested 48 hours post-transfection after fluorescent images were taken for subsequent experiments.

### RNA silencing assays using lentiviral transduction

To generate lentiviral particles, ∼12 × 10^6^ HEK293T packaging cells were seeded in complete DMEM in T175 flasks and incubated overnight. A transfection cocktail composed of packaging vectors [pMDLg/pRRE (3 μg; Addgene #12251), pRSV-Rev (1.5 μg; Addgene #12253), and pMD2.G/VSVg (1.8 μg; Addgene #12259)] and 14 μg of the target plasmid [RfxCas13d (Addgene #138149), or guide plasmid (cloned into Addgene #138150)] was combined with 150 μl of polyethylenimine (1 mg/ml) and topped up to 1.2-ml total volume with Opti-MEM. The solution was allowed to incubate for 20 min at RT, mixed well, and then added gently to packaging cells and incubated overnight at 37°C and 5% CO2. The following day media were refreshed, and for the next 48 hours, lentivirus-containing supernatants were harvested once daily. The harvested media were passed through a 0.4-um filter to remove cells and cellular debris and then concentrated via centrifugation for 15 min at 2000g in Amicon Ultra-15 (Merck) filter units. For target cell transduction, HEK293T cells were plated in a 12-well plate at a density of 1,000,000 cells per well (1mL). Polybrene (4 mg/ml) and the lentiviral supernatant diluted at various concentration for different desired MOI were added to the target cells and the cells were spun at 4000g for 1 hour. After two nights incubation at 37°C and 10% CO2, cells were split and DMEM media were refreshed. The transduced cells were then selected using blasticidin (10 μg/ml).

### Fluorescent microscopy analysis

For RNA silencing experiments, the fluorescence intensity was monitored using EVOS M5000 FL Cell Imaging System (Thermo Fisher). Images were taken 48 hours post-transfection for exogenous RNA targeting experiments, and 72 hours post-transfection for endogenous RNA targeting experiments. The fluorescence intensity of each image was quantified using a lab-written macro in ImageJ software. Briefly, all images obtained from a single experiment are simultaneously processed using a batch mode macro. First, images were converted to 8-bit, threshold adjusted, converted to black and white using the Convert to Mask function, and fluorescence intensity per pixel measured using Analyse Particles function. Each single mean fluorescence intensity was obtained from four different field of views for each crRNA and subsequently normalized to the non-targeting (NT) control crRNA. A two-fold or higher reduction in fluorescence intensity is considered biologically relevant.

### Cell flow cytometry

For β2M surface marker staining, up to 1×10^6^ cells were incubated in 50 µL of PBS/2% FBS (v/v) containing β2M antibody for 30 minutes on ice in the dark. The cells were then washed twice with 200 µL of PBS/2% (v/v) FBS before being re-suspended in 200 µL of PBS/2% FBS (v/v) for flow cytometry analysis. Flow cytometry analysis was performed using either the FACS Symphony Cell Analyser A5 or A3 (BD Biosciences). All flow cytometry profiles were analysed using FlowJo V10 software (Tree Star Inc). Supplementary Table 2 lists the antibodies that were used for cell flow cytometry.

### Western Blot

Cells were washed three times with ice-cold PBS -/- and lysed on ice in RIPA lysis buffer [50 mM Tris (Sigma-Aldrich, T1530), pH 8.0, 150 mM NaCl, 1% NP-40 (Sigma-Aldrich, I18896), 0.1% SDS, 0.5% sodium deoxycholate (Sigma-Aldrich, D6750)] containing protease inhibitor cocktail (Roche, 04693159001) and phosphatase inhibitor cocktail (Roche, 4906845001). Samples were incubated for 30min at 4 °C with rotation (25 rpm), and centrifuged at 16,000 g for 10 min, 4 °C. Supernatant was transferred to a new tube. Protein concentrations were quantified using the Pierce BCA Protein Assay Kit (Thermo Fisher, 23225) according to the manufacturer’s instructions. A total of 10 μg of protein diluted in 1x Bolt LDS sample buffer (Thermo Fisher, B007) and 1x Bolt sample reducing agent (Thermo Fisher, B009) were denatured at 95 °C for 5 min. Samples were resolved by Bolt Bis-Tris Plus 4–12% gels (Thermo Fisher, NW04120BOX) in 1x MES SDS running buffer (Thermo Fisher, B0002) and transferred to 0.45 μM PVDF membranes (Thermo Fisher, 88518) by a Trans-Blot Semi-Dry electrophoretic transfer cell (Bio-Rad) at 20 Volt for 30 min. Membranes were incubated in blocking buffer 5% (w/v) BSA (Sigma-Aldrich, A3059) in TBST with 0.15% Tween 20 (Sigma-Aldrich, P1379) for 1h at RT and probed overnight with primary antibodies at 4 °C. Blots were washed three times in TBST with 0.15% Tween20, followed by incubation with fluorophore-conjugated or HRP-conjugated secondary antibodies for 1h at RT. Membranes were washed in TBST (0.15% Tween20) three times and chemiluminescence was detected using the ChemiDoc Imaging System (Bio-Rad). The antibodies used for western blots are listed in Supplementary Table 2.

### RNA extraction, RNA sequencing and RT-PCR

Total RNA was isolated by standard Trizol-chloroform extraction according to manufacturer’s instruction (Thermo Fisher, 15596026), followed by DNase treatment with RQ1 RNase-Free DNase according to manufacturer’s instruction (Promega, M6101). RNA integrity was analysed by Agilent 2200 Tapestation (Agilent, G2964AA) with RNA ScreenTape (Agilent, 5067- 5576) according to the manufacturer’s instructions.

For zebrafish RT-PCR, mRNA was first isolated from 10 embryos of either Tg(fli1a:EGFP) or Tg(kdrl:EGFP) transgenics, or embryos injected with 200 pg of rEGFP mRNA (n=6) at 2 dpf using the Direct-zol RNA kit (Zymo Research, R2050) according to manufacturer’s instructions. 1.7 μg total RNA was used to synthesise cDNA using the high-capacity cDNA reverse transcription kit (Thermo Fisher, 4368814) following the manufacturer’s instructions. Quantitative RT-PCR reaction was performed in technical triplicate using the Applied Biosystems QuantStudio 7 Flex (Thermo Fisher) and the PowerUp™ SYBR™ Green Master Mix (Thermo Fisher, A25742). The reaction contained 4 μL of cDNA (1/15 dilution), 0.5 μL of forward primer, and 0.5 μL of reverse primer, and ribosomal protein 13L (rp113a) and eukaryotic translation elongation factor 1 alpha 1, like 1 (eef1a111) were used as reference genes. Primer sequences are detailed in Supplementary Table 3 Data was analysed as previously published (PMID 11846609).

RNA was processed by Australian Genome Research Facility (AGRF) using Illumina Ribo-Zero Plus rRNA Depletion Kit and pair-end sequenced on an Illumina NextSeq 500. Libraries (triplicate RNA samples) were created from the samples and combined for sequencing with 150bp reads, for a total of ∼50 million reads per HEK293T cell sample.

### Mass spectrometry proteomics analysis

At 48h post-transfection, cells were washed three times with ice-cold PBS -/- and lysed on ice in RIPA lysis buffer [50 mM Tris (Sigma-Aldrich, T1530), pH 8.0, 150 mM NaCl, 1% NP-40 (Sigma-Aldrich, I18896), 0.1% SDS, 0.5% sodium deoxycholate (Sigma-Aldrich, D6750)] containing protease inhibitor cocktail (Roche, 04693159001) and phosphatase inhibitor cocktail (Roche, 4906845001). 10 mM of TCEP (Tris(2-carboxyethyl)phosphine) were added to 50 µg of protein lysates and incubated at 65 °C for 20 min for disulfide bonds reduction. Iodoacetamide was added to a final concentration of 55mM to alkylate proteins, samples were incubated at 37°C for 45 minutes in the dark. Samples were acidified by the addition of 2.5% final concentration of phosphoric acid. Then, S-Trap binding buffer (90% MeOH, 100 mM Triethylammonium bicarbonate (TEAB), pH 7.1) was added to the acidified lysate, and samples were passed through Suspension trapping (S-trap) column. The bound samples were then washed three times with the S-Trap binding buffer. The bound proteins were then digested overnight at 37°C by trypsin in 50 mM TEAB at 1:10 ratio. The peptides were eluted with 40 µL of 50 mM TEAB, followed by 40 µL 0.2% aqueous formic acid, and finally with 35 µL of 50% acetonitrile containing 0.2% formic acid. Samples were freeze dried overnight in a freeze dryer (Eppendorf Concentrator Plus), resuspended in 20 µL 2% acetonitrile, 0.1% trifluoroacetic acid, sonicated for 10 min at room temperature, centrifuged at 20,000 × g for 10 min, and transferred into HPLC vials for analysis.

Samples were analysed by LC-MS/MS using Orbitrap Exploris 480 (Thermo Scientific) fitted with nanoflow reversed-phase-HPLC (Ultimate 3000 RSLC, Dionex). The nano-LC system was equipped with an Acclaim Pepmap nano-trap column (Dionex – C18, 100 Å, 75 μm × 2 cm) and an Acclaim Pepmap RSLC analytical column (Dionex – C18, 100 Å, 75 μm × 50 cm). Typically for each LC-MS/MS run, 1 μL of the peptide mix was loaded onto the enrichment (trap) column at an isocratic flow of 5 μL/min of 2% Acetonitrile containing 0.05% trifluoroacetic acid for 6 min before the analytical column is switched in-line. The buffers used for the LC were 0.1% v/v formic acid in water (solvent A) and 100% Acetonitrile/0.1% formic acid v/v (Solvent B). The gradient used was 3% B to 23% B for 89 min, 23% B to 40% B in 10 min, 40% B to 80% B in 5 min and maintained at 80% B for the final 5 min before equilibration for 10 min at 2% B prior to the next analysis. All spectra were acquired in positive mode with full scan MS spectra scanning from m/z 300-1600 at 120,000 resolutions with AGC target of 3×10^6^ with a maximum accumulation time of 25 ms. The peptide ions with charge state ≥2-6 were isolated with an isolation window of 1.2 m/z and fragmented with a normalized collision energy of 30 at 15,000 resolution.

All generated files (except for RNA transfection PspCas13b target titration experiment) were analysed with MaxQuant (version 2.2.0.0) and its implemented Andromeda search engine to obtain protein identifications as well as their label-free quantitation (LFQ) intensities. Database searching was performed with the following parameters: cysteine carbamidomethyl as a fixed modification; oxidation and acetyl as variable modifications with up to 2 missed cleavages permitted; main mass tolerance of 4.5 ppm; 1% protein false discovery rate (FDR) for protein and peptide identification; and minimum two peptides for pair-wise comparison in each protein for label-free quantitation. The raw data files were searched against the modified Human reference proteomes (UP000005640) supplemented with the amino acid sequence of the exogenous proteins mCherry, EGFP, LwaCas13a, PspCas13b, and RfxCas13d.

Mass spectrometry data from the PspCas13b target-titration experiment (RNA samples) were analysed using Dia.NN (version 1.9.1) [57]. Dia.NN was run with modified parameters relative to the default settings, allowing up to two missed cleavages and one variable modification per peptide. Oxidation of methionine (Ox[M]) and N-terminal acetylation (Ac[N-term]) were included as variable modifications. The neural-network classifier was set to double-pass mode, and a 1% false discovery rate (FDR) was applied at both the protein and peptide levels for identification.

The MaxQuant and Dia.NN result output was further processed with Perseus (Version 2.0.7.0), a module from the MaxQuant suite. After removing reversed and known contaminant proteins, the LFQ values were log_2_ transformed and the reproducibility across the biological replicates was evaluated by a Pearson’s correlation analysis. The replicates were grouped accordingly, and all proteins were removed that had less than 4 “valid value” in each group. The missing values were replaced by imputation, and differential expression analysis between targeting and non-targeting samples was performed using two-sample t test. Proteins with log_2_ fold change (FC)>1 were considered as upregulated while proteins with log_2_FC<-1 were considered downregulated. Adjusted p-value (with 1% false discovery rate FDR) < 0.05 was considered as statistically significant. Genes that were significantly up or down regulated between targeting and NT crRNAs are highlighted in the volcano and linear regression plots. All proteomics data are provided with the paper in a source data file.

### Resazurin assay

A resazurin sodium salt working solution (Sigma-Aldrich, R7017-1G) was prepared by resuspension in PBS (pH 7.4) to a final concentration of 0.15 mg/mL. Working solutions were filter sterilized and stored at 4°C in light-protected tubes. At each timepoint, 100uL of resazurin working solution is added to each of the conditions in a 24-well plate and incubated at 37°C for 2-3 hours. Then each well was pipetted into 4 or 5 technical replicates (100 μL each) into a 96-well plate for absorbance measurement. Resazurin reduction was calculated by subtracting the background absorbance at 600 nm from the absorbance at 570 nm using the Cytation 5 Multimode Plate Reader (Agilent), with data normalized to the NT crRNA conditions.

### CellTiter-Glo luminescence assay

CellTiter-Glo buffer (Promega, G756A) was combined with the CellTiter-Glo substrate (Promega, G755A) according to the manufacturer’s instructions to prepare the working solution. Culture media were removed from cells in 24-well plates, and 500 µL of the CellTiter-Glo working solution was added to each well. Plates were incubated for 10 min at room temperature to allow cell lysis and luminescence development. Subsequently, 100 µL from each well was transferred into three technical replicates in a 96-well opaque plate (Merck, MSSBNFX40). Luminescence was measured using a Cytation 5 Multimode Plate Reader (Agilent). Luminescence values were normalized to the non-targeting (NT) crRNA control condition.

### Zombie NIR viability staining

Cells cultured in 24-well plates were washed once with protein-free PBS and resuspended in 100 µL of diluted Zombie NIR viability dye (BioLegend, 420201; 1:500 in PBS). Cells were incubated for 15–30 min at room temperature in the dark, followed by one wash with PBS containing 1–2% BSA or serum. Stained cells were then resuspended in staining buffer and subjected to flow cytometry analysis.

### Zebrafish maintenance and Imaging

All zebrafish experiments and protocols were approved by and carried out in compliance to guidelines of the animal ethics committees at La Trobe University (AEC24019). The transgenic zebrafish strains Tg(fli1a:nEGFP)^y7^ [47], Tg(kdrl:EGFP)^s843^ (PMID: 16107477), Tg(kdrl:HRAS-mCherry)^s916^ (Tg(kdrl:mCherry), PMID 19287381), and Tg(elavl3:EGFP)^knu3^ (PMID: 11071755) were used in this study. Embryos were either dechorionated, anaesthetised in 0.08 mg/mL tricaine and mounted (1 and 2 dpf), or mounted with chorion (4 hpf) as previously described [58]. Zebrafish live-imaging was conducted using either a Nikon SMZ800N microscope (with attached TrueChromo 4K Pro camera for bright field images), an Andor Dragonfly 202 spinning disc confocal microscope, or a Nikon SMZ18 fluorescent microscope. To obtain bright field videos, unanaesthetised 3 dpf larvae were collected into middle of the camera view (SMZ800N microscope with attached TrueChromo 4K Pro camera) and touched using a superfine eyelash brush No. 1 (ProSciTech, Australia).

### Zebrafish microinjections

RfxCas13d-based knockdowns in zebrafish were conducted as previously described (PMID 35005640) with modifications. Briefly, wild-type (AB) single-cell-staged embryos were injected with 12.5, 25, 50 or 200 pg of EGFP mRNA (concentrations 12.5, 25, 50 or 200 ng/μL), generated using the mMessage mMachine SP6 Transcription Kit (Thermo Fisher Scientific) following manufacturer’s instructions, 250 pg RfxCas13d-mFlame mRNA (concentration 250 ng/μL), and 250 pg of either EGFP-targeting crRNA or non-targeting crRNA (concentration 250 ng/μL) directly into the cell. Alternatively, single-cell-staged embryos from outcrosses between heterozygous transgenic adult (Tg(fli1a:EGFP), Tg(kdrl:EGFP), Tg(kdrl:mCherry), or Tg(elavl3:EGFP)), and wild type (AB), as indicated, were injected with 250 pg RfxCas13d-mFlame mRNA, and 250 pg of either EGFP-targeting crRNA or non-targeting crRNA directly into the cell.

### Image processing and analysis

All spinning disc confocal Images were processed with image processing software FIJI (PMID: 22743772). ISV height (horizontal height from the base ISV/dorsal aorta intersection to the tip of the ISV) was measured using FIJI. The number of fully formed ISVs (ISVs that have reached the level of the DLAV) or DLAV (fully connected DLAV) between the 10^th^ to the 14^th^ somites (from the anterior side, 6 ISVs) were manually counted on FIJI and represented as ratio per embryo. Fish movement upon touching was quantified using a numerical range between 0 and 1, where 0 meant that fish didn’t move while 1 meant that fish exited the camera’s field of view. Ranges in between (0.125, 0.25, 0.5, and 0.75) were variations of these distances (e.g., 0.75 represents fish that have moved 75% of the way towards the end of the camera’s view).

## Data analysis

Data analyses and visualizations (graphs) were performed in GraphPad Prism software version 9 and R Studio. The silencing efficiency of various crRNAs was analysed using unpaired two-tailed Student’s t-test where we compare every mean to a control mean as indicated in the Figures (95% confidence interval). The p values are indicated in the Figures. p < 0.05 is considered statistically significant.

Supplementary table 1 – crRNA, targets and plasmids

Supplementary table 2 – antibody

Supplementary table 3 - primers

## Funding

This work was supported by a Cancer Council Victoria Ventures grant (829606); mAP mRNA Victoria grant (RCH0153742 to M.F), the Cummings Centre grant (051848 to M.F, S.R.L and W.Z), and the La Trobe University ABC Internal Investment Scheme (to KSO). HC and VI were supported by a PhD scholarship from the University of Melbourne and La Trobe University, respectively. RWJ was supported by the National Health and Medical Research Council (NHMRC) and the Barrie Dalgleish Centre for Myeloma and associated Blood Diseases. The study was generously supported by The Peter MacCallum Cancer Foundation.

## Acknowledgments

The authors thank all lab members from the Fareh, Trapani, and Okda lab for facilitating experiments and discussions. We also thank Prof. Ilia Voskoboinik for the valuable discussions and critical feedback.

## Authors contributions

MF and HC conceived the study. MF, KSO, and JAT supervised the study. HC and MF designed the human cell experiments. HC performed the experiments in human cells and analysed the data assisted by WH, GJS, JK, CS, and PR. All Zebrafish experiments were performed in the lab of KSO by VI and SB with input from KSO, HC, and MF. SP helped optimise initial RNA injection in Zebrafish embryos under the supervision of BMH. WH, VI, GJS, JK, CS, PR, JMLC, WZ, SRL, RWJ, BMH, SJV, IV, and JAT discussed the data and provided critical input regarding mRNA design and collateral activity in human cells and zebrafish embryos. HC generated the figures with input from MF and KSO. HC, KSO, and MF wrote the manuscript. All authors read, commented on, edited and approved the manuscript.

## Competing interests

The Johnstone lab receives research funding from Pfizer, BMS, MycRx, and AstraZeneca. R.W.J. is a co-founder and shareholder of MycRx and receives consultancy payments. The other authors declare no conflict of interest.

