## Supplementary material for "Target RNA abundance controls the collateral activity of RfxCas13d in human cells and zebrafish embryos": Suppl. File

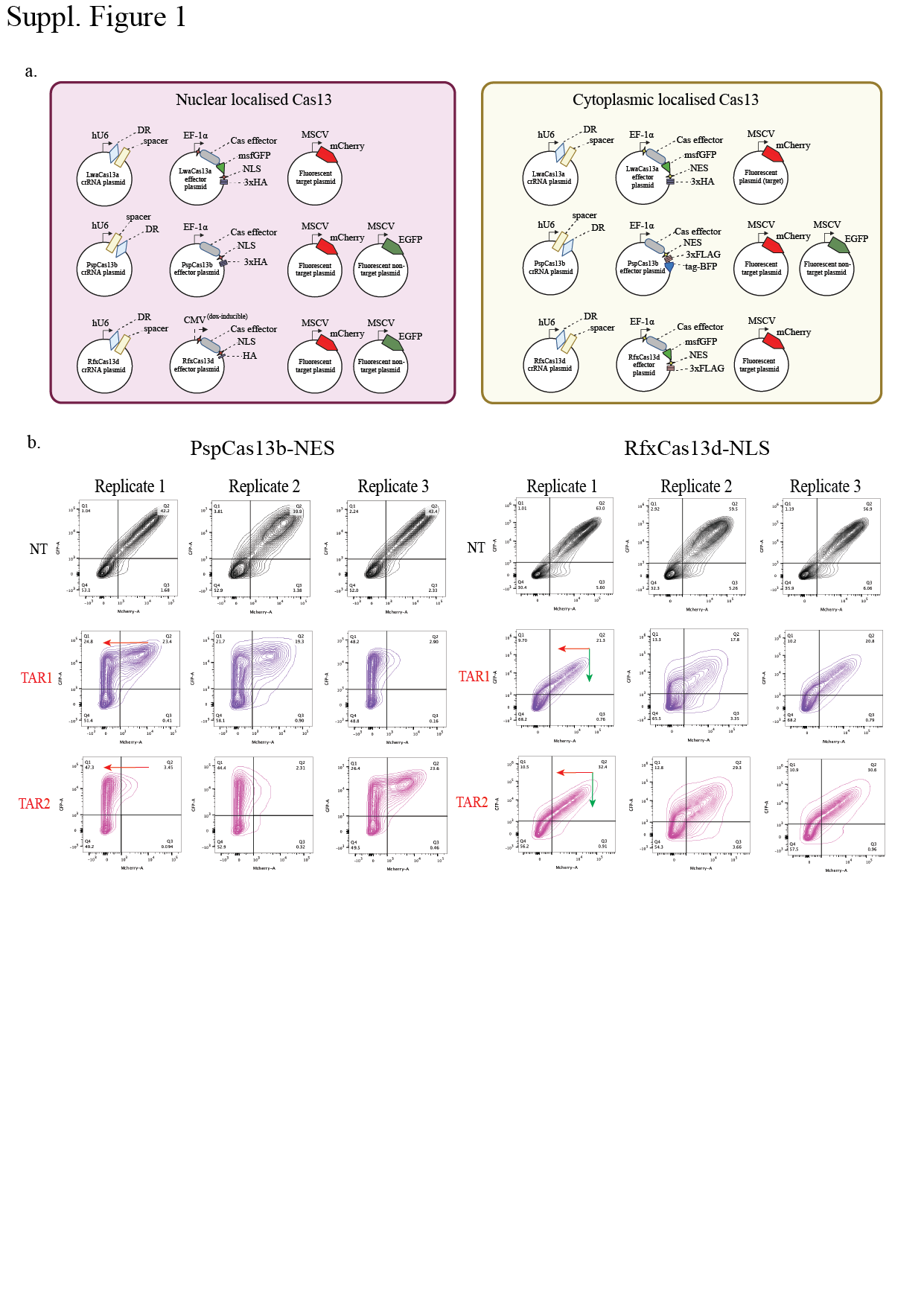
**Suppl. Figure 1**

**a,** Schematics of the three differently localised Cas13 orthologues, crRNA, mCherry and EGFP plasmid constructs used for the silencing assay. **b**, Silencing of mCherry and EGFP by PspCas13b (NES) or RfxCas13d (NLS) shown as contour plots. The x-axis shows mCherry (target) silencing and the y-axis shows EGFP (non-target) silencing at 48 hours post transfection in HEK293T cells. Black graphs represent non-targeting (NT) crRNA, purple and magenta graphs represent the two mCherry-targeting crRNA conditions (n=3).

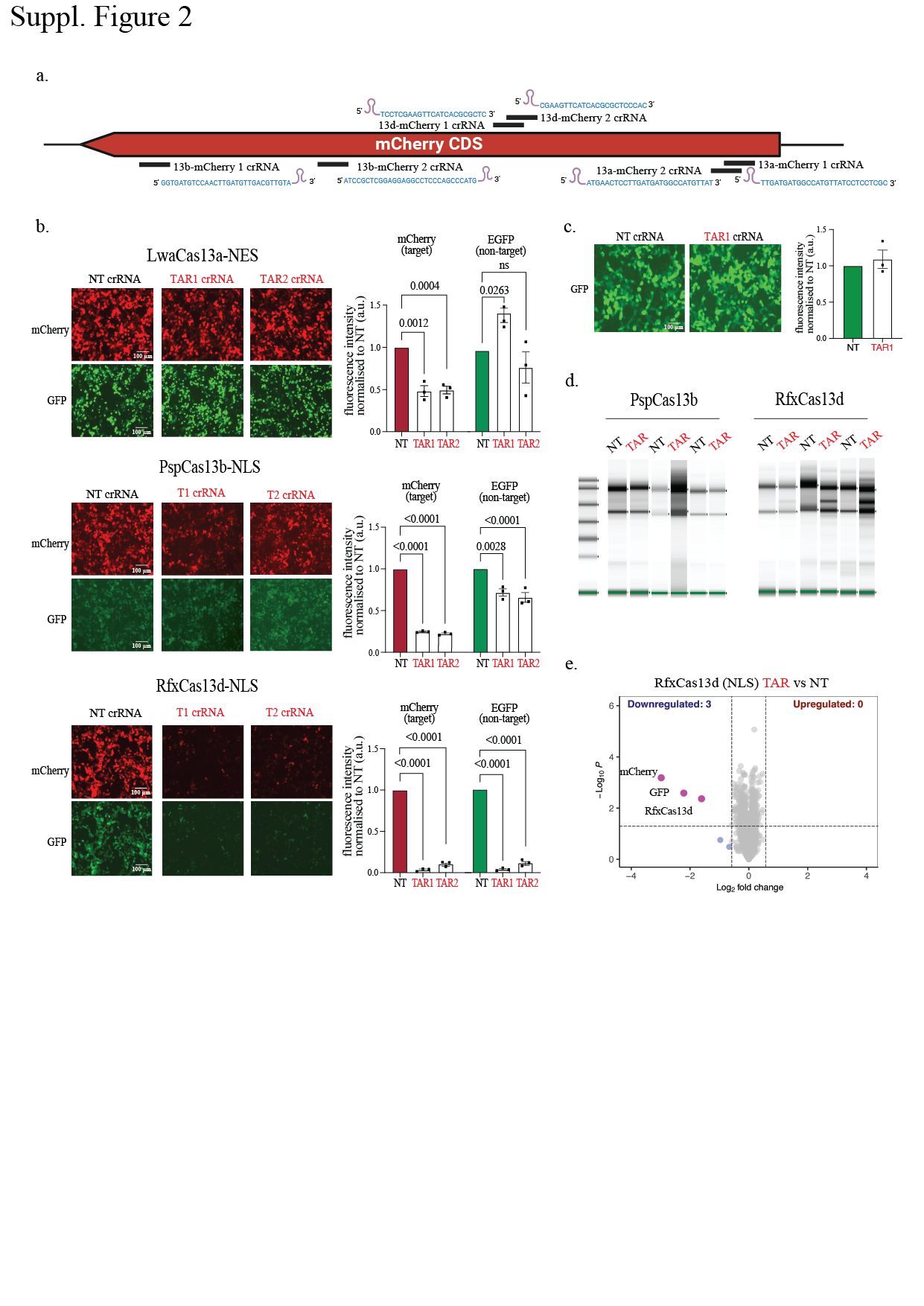

**Suppl. Figure 2**

**a**, Schematic of crRNA sequence designs matching different regions of the target mCherry RNA for either LwaCas13a, PspCas13b or RfxCas13d. **b**, **Left**, Representative fluorescence microscopy images of the silencing of mCherry and EGFP RNA by the LwaCas13a (NES), PspCas13b (NLS), and RfxCas13d (NLS) ortholog with two mCherry-targeting crRNAs versus a non-targeting (NT) control crRNA in HEK293T cells (n = 3). Scale bar is 100μm; **Right**, Quantification of Cas13 silencing of EGFP transcripts with either a NT or an mCherry-targeting crRNA in the absence of mCherry target RNA. Data points in the graph are averages of the normalized mean fluorescence from four representative fields of view imaged (n = 3). The data are represented in arbitrary units (a.u.). Errors are the s.e.m. and P values of Student’s t-test are indicated (95% confidence interval). **c**, **Left**, Representative fluorescence microscopy images of the silencing of EGFP (non-target) without mCherry target transcripts by RfxCas13d ortholog with a potent mCherry-targeting crRNAs versus a non-targeting (NT) control crRNA in HEK293T cells (n = 3); **Right**, Quantification of Cas13 silencing of EGFP transcripts with either NT or mCherry-targeting crRNA. Data points in the graph are averages of the normalized mean fluorescence from four representative fields of view imaged (n = 3). The data are represented in arbitrary units (a.u.). Errors are the s.e.m. and p values of Student’s t-test are indicated (95% confidence interval). **d,** TapeStation showing 28S and 18S ribosomal RNA of the RNA extracts from HEK293T cells transfected with PspCas13b or RfxCas13d with mCherry and NT or mCherry-targeting crRNA. RfxCas13d with mCherry-targeting crRNA shows additional third band of 28s rRNA cleavage, indicating the presence of collateral cleavage. **e,** Volcano plots showing the proteome of HEK293T cells expressing mCherry, EGFP, mCherry-targeting or NT crRNA paired with RfxCas13d (NLS). Data points represent proteins, with log_2_FC > 0.58 (upregulation, blue) or log_2_FC < −0.58 (downregulation, red) and p < 0.05 indicating significantly differential expression. Total downregulated and upregulated proteins are shown at the top corners. Exogenous Cas13, mCherry and EGFP are labelled purple. Results were analysed by an unpaired two-tailed Student’s t-test (n = 5).

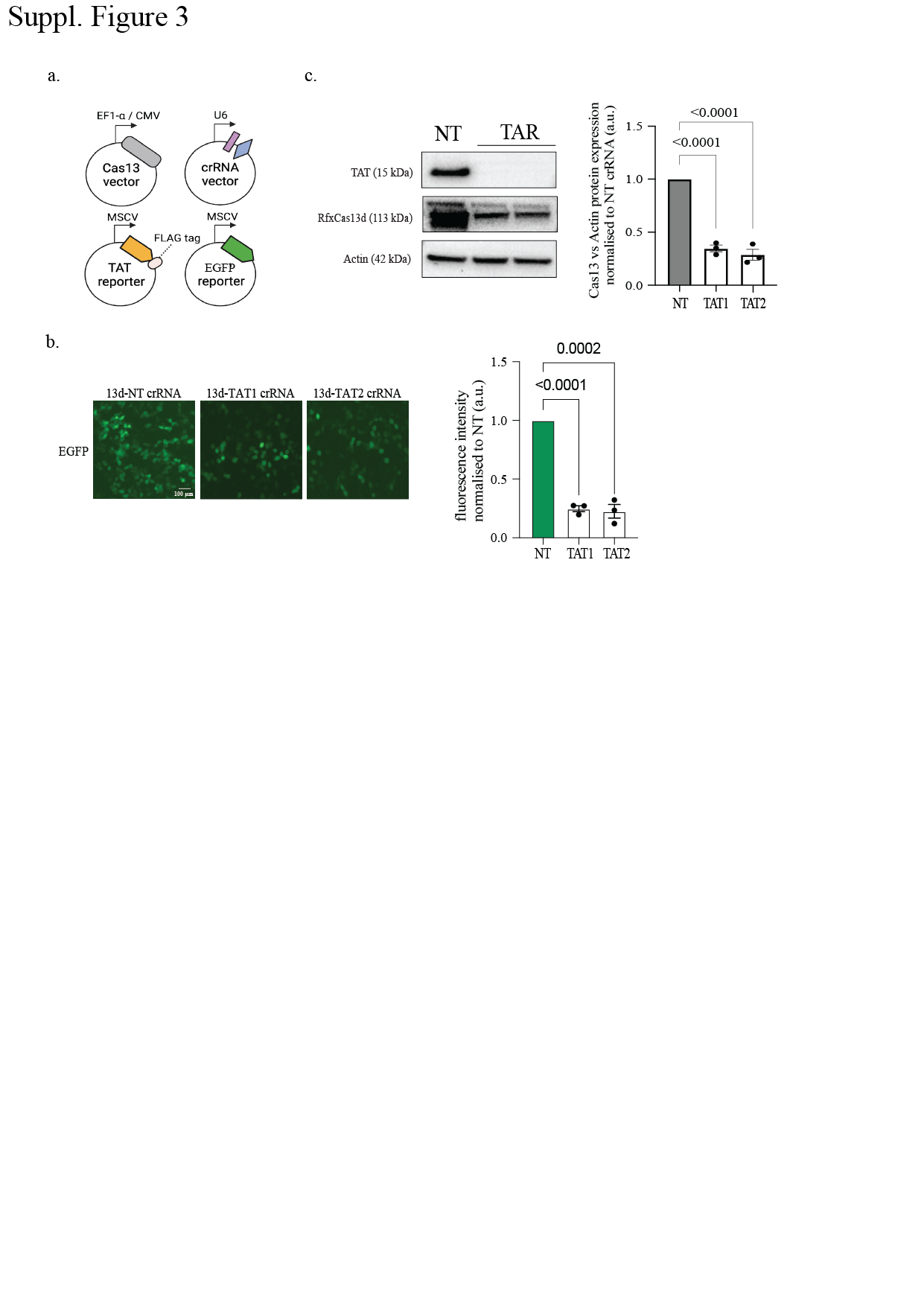

**Suppl. Figure 3**

**a,** Schematic of RfxCas13d plasmid transfection silencing assay to assess exogenously introduced TAT on-target cleavage and EGFP collateral cleavage. **b,** **Left,** Representative fluorescence microscopy images of the silencing of EGFP (non-target) by the RfxCas13d ortholog with a TAT-targeting crRNA versus a non-targeting (NT) control crRNA in HEK293T cells (n = 3); **Right,** Quantification of Cas13 silencing of EGFP transcripts with either NT or TAT-targeting crRNA. Data points in the graph are averages of the normalized mean fluorescence from four representative fields of view imaged (n = 3). The data are represented in arbitrary units (a.u.). Errors are the s.e.m. and p values of Student’s t-test are indicated (95% confidence interval). **c, Left,** Representative Western blot of RfxCas13d, TAT and β actin protein levels in HEK 293T cells expressing either TAT-targeting crRNA or NT crRNA. **Right,** Quantification of Cas13 protein silencing normalised to β actin protein levels (n = 3). The data are represented in arbitrary units (a.u.). Errors are the s.e.m. and p values of Student’s t-test are indicated (95% confidence interval).

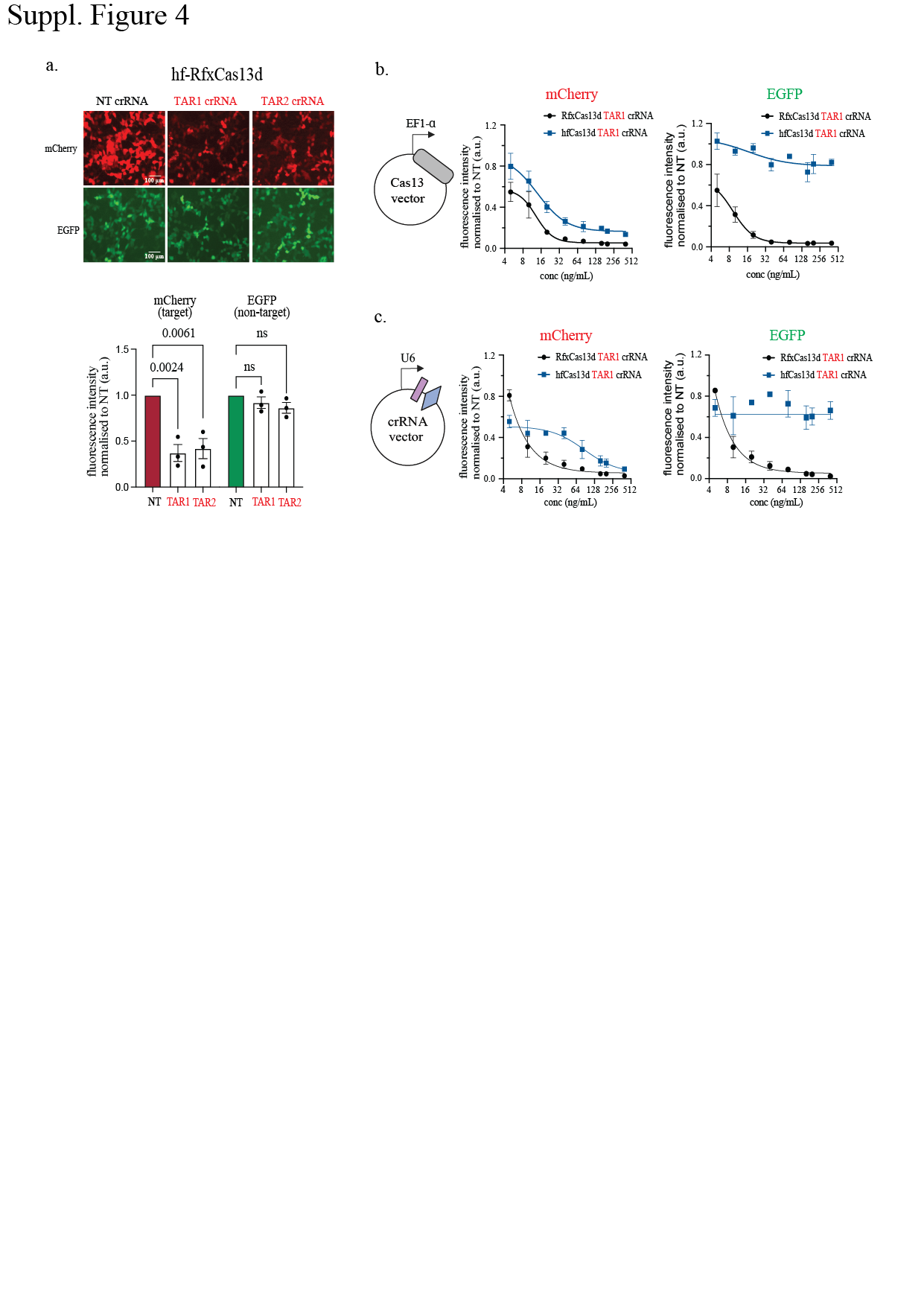
**Suppl. Figure 4**

**a,** Schematic of synthetic hfCas13d effector protein with 4 amino acid mutations from RfxCas13d, which supposedly eliminates collateral cleavage without affecting on-target cleavage. Scale bar is 100μm. **b**, **Top,** Representative fluorescence microscopy images of the silencing of mCherry and EGFP transcripts by hfCas13d with two potent mCherry-targeting crRNAs versus a non-targeting (NT) control crRNA in HEK293T cells (n = 3); **Bottom,** Quantification of Cas13 silencing of mCherry and EGFP transcripts with either NT or mCherry-targeting crRNAs. Data points in the graph are averages of the normalized mean fluorescence from four representative fields of view imaged (n = 3). The data are represented in arbitrary units (a.u.). Error bars are the s.e.m. and P values of Student’s t-test are indicated (95% confidence interval). **c&d,** Quantification of the comparison of RfxCas13d (black) and hfCas13d (blue) silencing of mCherry and EGFP transcripts with a titration of either the Cas13 effector proteins or the crRNAs. Data points in the graph are averages of the normalized mean fluorescence from four representative fields of view imaged (n = 3).

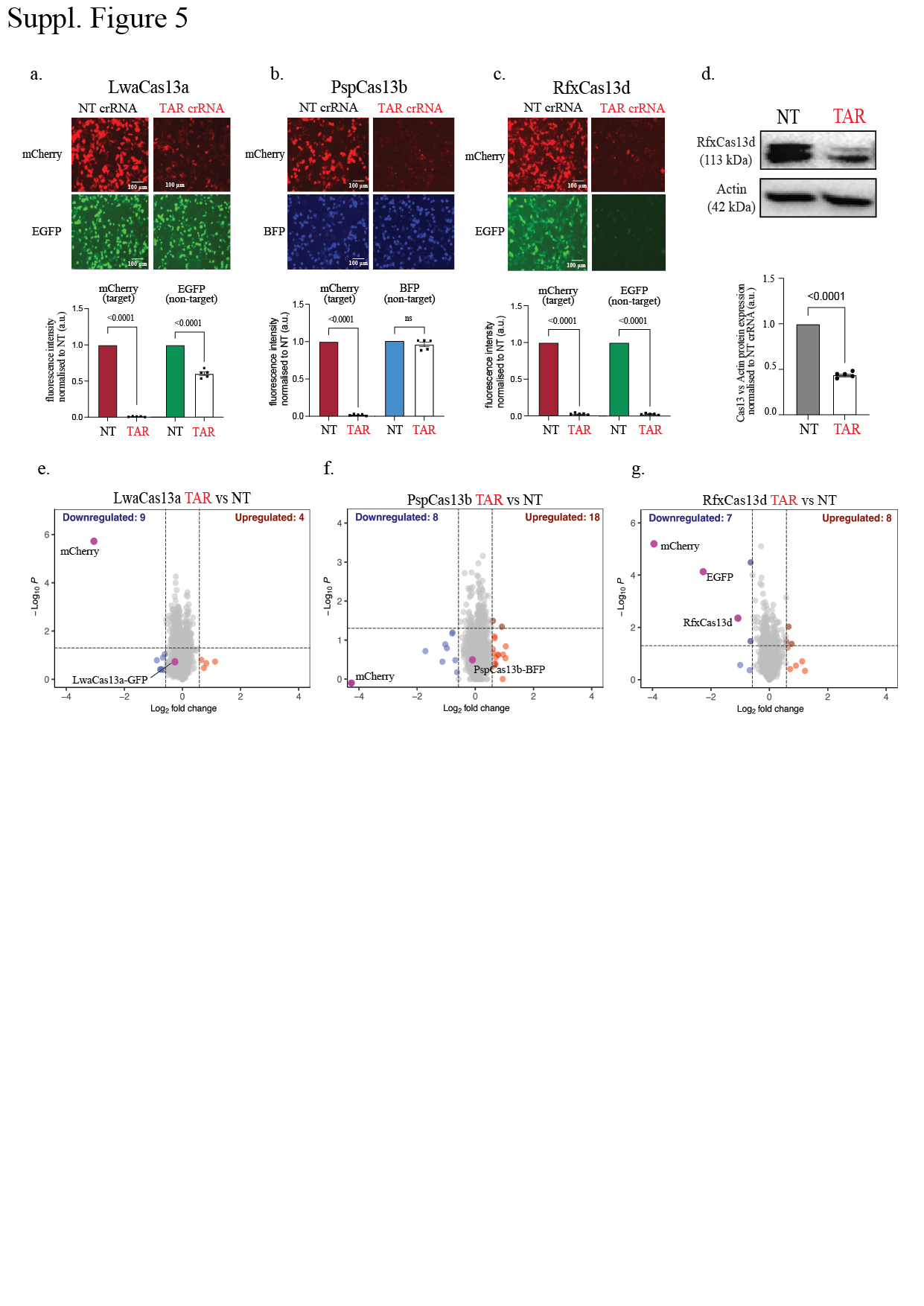

**Suppl. Figure 5**

**a-c, Top,** Representative fluorescence microscopy images of the silencing of mCherry and EGFP transcripts by three Cas13 orthologs with two potent mCherry-targeting crRNAs versus a non-targeting (NT) control crRNA in HCT116 cells (n = 3); **Bottom**, Quantification of Cas13 silencing of mCherry (on-target cleavage) and EGFP (collateral cleavage) transcripts with mCherry-targeting crRNA compared to NT crRNA. Data points in the graph are averages of the normalized mean fluorescence from four representative fields of view imaged (n = 3). The data are represented in arbitrary units (a.u.). Errors are the s.e.m. and p values of Student’s t-test are indicated (95% confidence interval). Scale bar is 100μm. **d,** Representative Western blot and quantification of the expression level of RfxCas13d ortholog effector protein levels in HCT116 cells expressing either mCherry-targeting crRNA or NT crRNA normalised to β actin (n = 3). **e-g,** Volcano plots showing the proteome of cells expressing mCherry, EGFP, mCherry-targeting or NT crRNA paired with different Cas13 orthologs in HCT116 cells. Data points represent proteins, with log_2_FC > 0.58 (upregulation, blue) or log_2_FC < −0.58 (downregulation, red) and P < 0.05 indicating significantly differential expression. Total downregulated and upregulated proteins are shown at the top corners. Exogenous Cas13, mCherry, and EGFP are labelled purple. Results were analysed by an unpaired two-tailed Student’s t-test (n = 5).

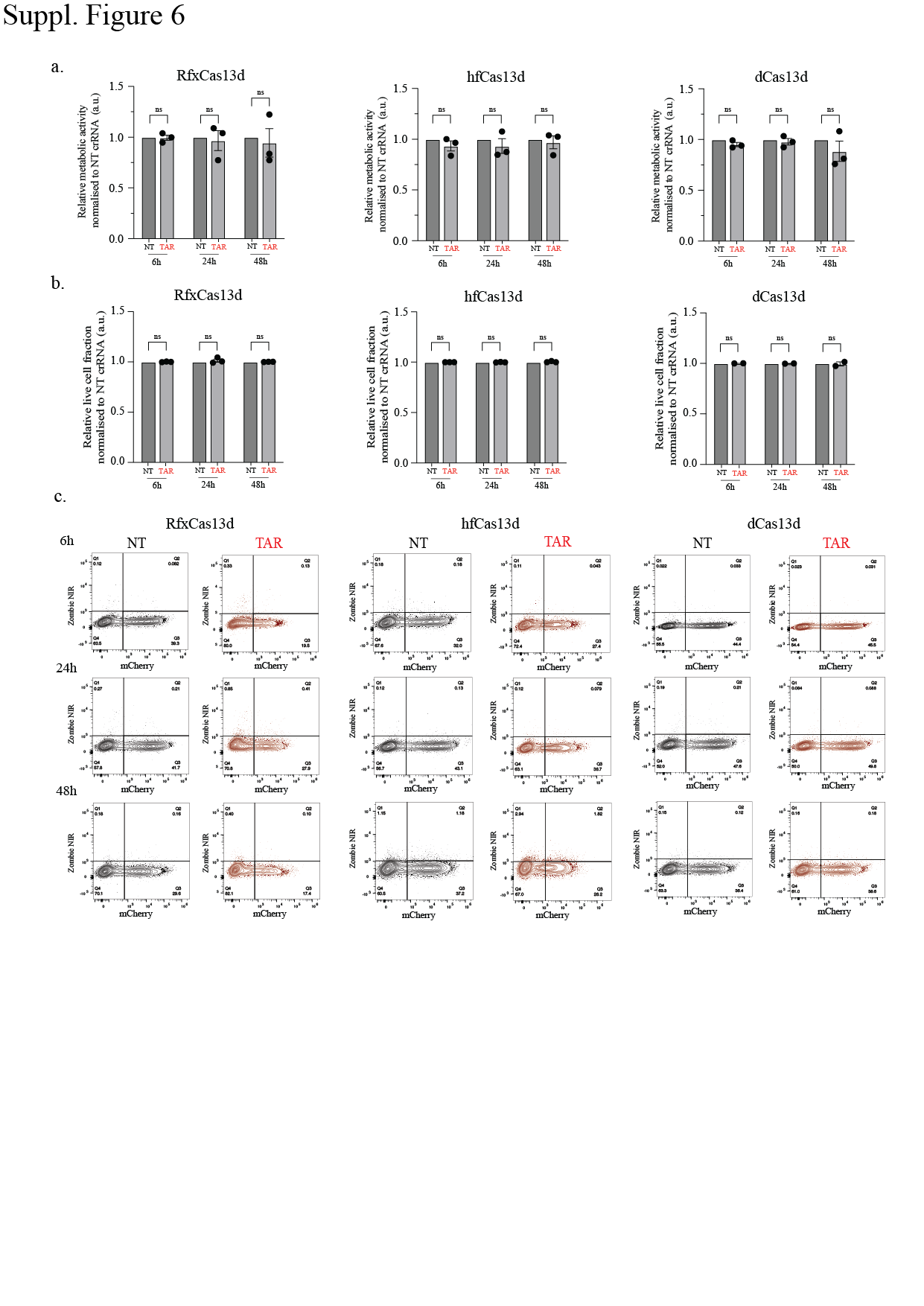

**Suppl. Figure 6**

**a,** Resazurin cell metabolism assay performed for HEK 293T cells transfected with mCherry, EGFP, mCherry-targeting or non-targeting (NT) crRNA paired with either RfxCas13d or hfCas13d at 24, 48, and 72 hours respectively. Data points in the graph are averages of the normalized to the NT crRNA conditions for each Cas13 effector protein (n = 3). Error bars are the s.e.m.. **b,** Quantification of percentage of cells alive from the Zombie Near Infrared (ZIR) cell viability assay performed for HEK293T cells transfected with mCherry, EGFP, mCherry-targeting (coloured) or NT crRNA (black) paired with either RfxCas13d or hfCas13d at 24, 48, and 72 hours respectively. Data points in the graph are averages of the normalized to the NT crRNA conditions for each Cas13 effector protein (n = 3). **c,** FACS plot of ZIR assay showing representative images of the silencing of mCherry as well as dead cell positively stained with ZIR dye.

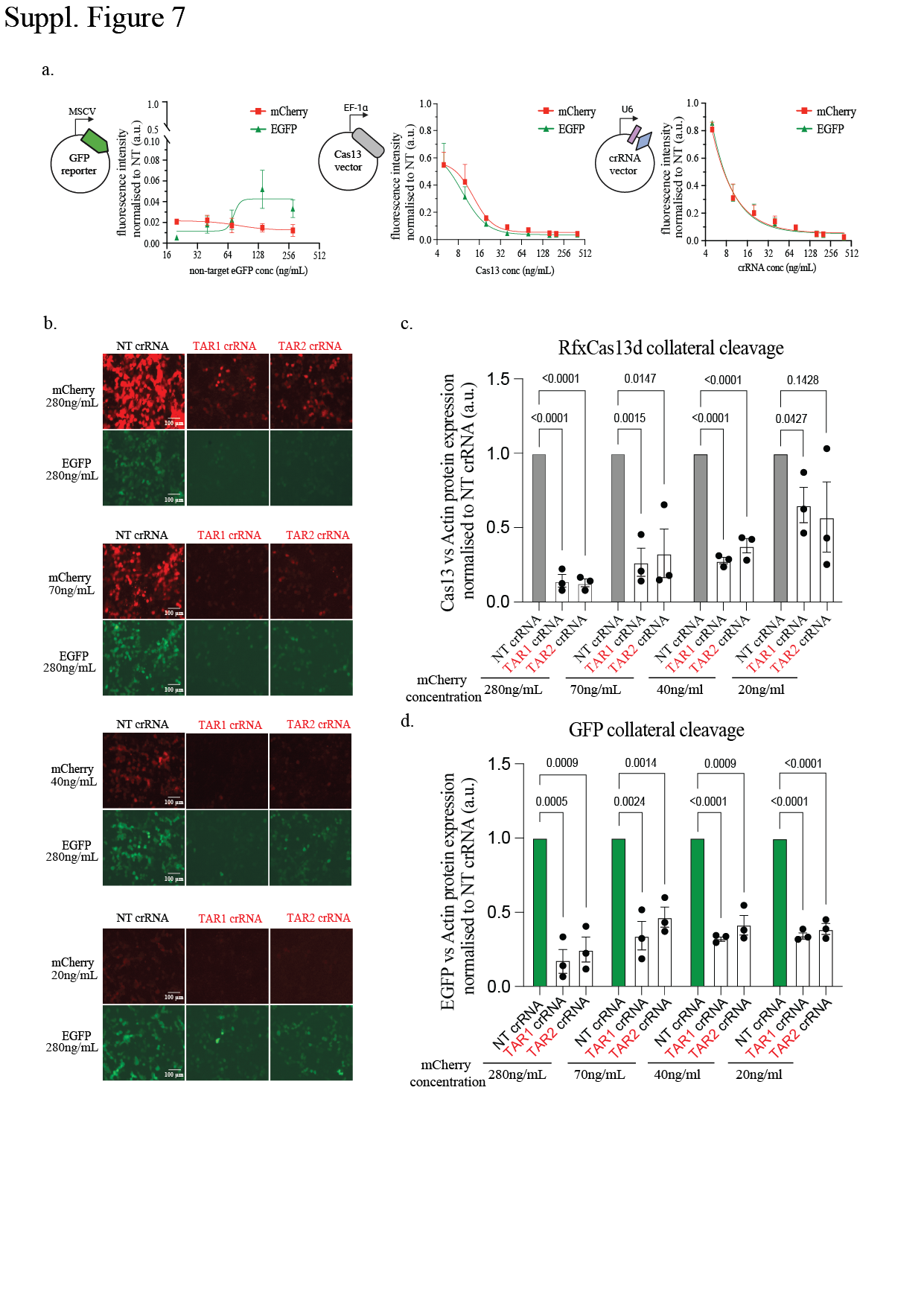

**Suppl. Figure 7**

**a,** Quantification of the comparison of on-target (mCherry) and collateral cleavage (EGFP) silencing with a titration of either of the three plasmid transfection components: Cas13 effector protein, NT or mCherry-targeting crRNA, and EGFP non-target. Data points in the graph are averages of the normalized mean fluorescence from four representative fields of view imaged (n = 3). **b,** Representative fluorescence microscopy images of the silencing of mCherry and EGFP transcripts by RfxCas13d with two potent mCherry-targeting crRNAs versus a non-targeting (NT) control crRNA in HEK293T cells at the highest mCherry concentration (280 ng/mL) and the lowest mCherry concentration (20 ng/mL). Scale bar is 100μm. **c&d**, Quantification of RfxCas13d and EGFP protein silencing normalised to β actin protein levels (n = 3 for each concentration). The data are represented in arbitrary units (a.u.). Errors are the s.e.m. and p values of Student’s t-test are indicated (95% confidence interval).

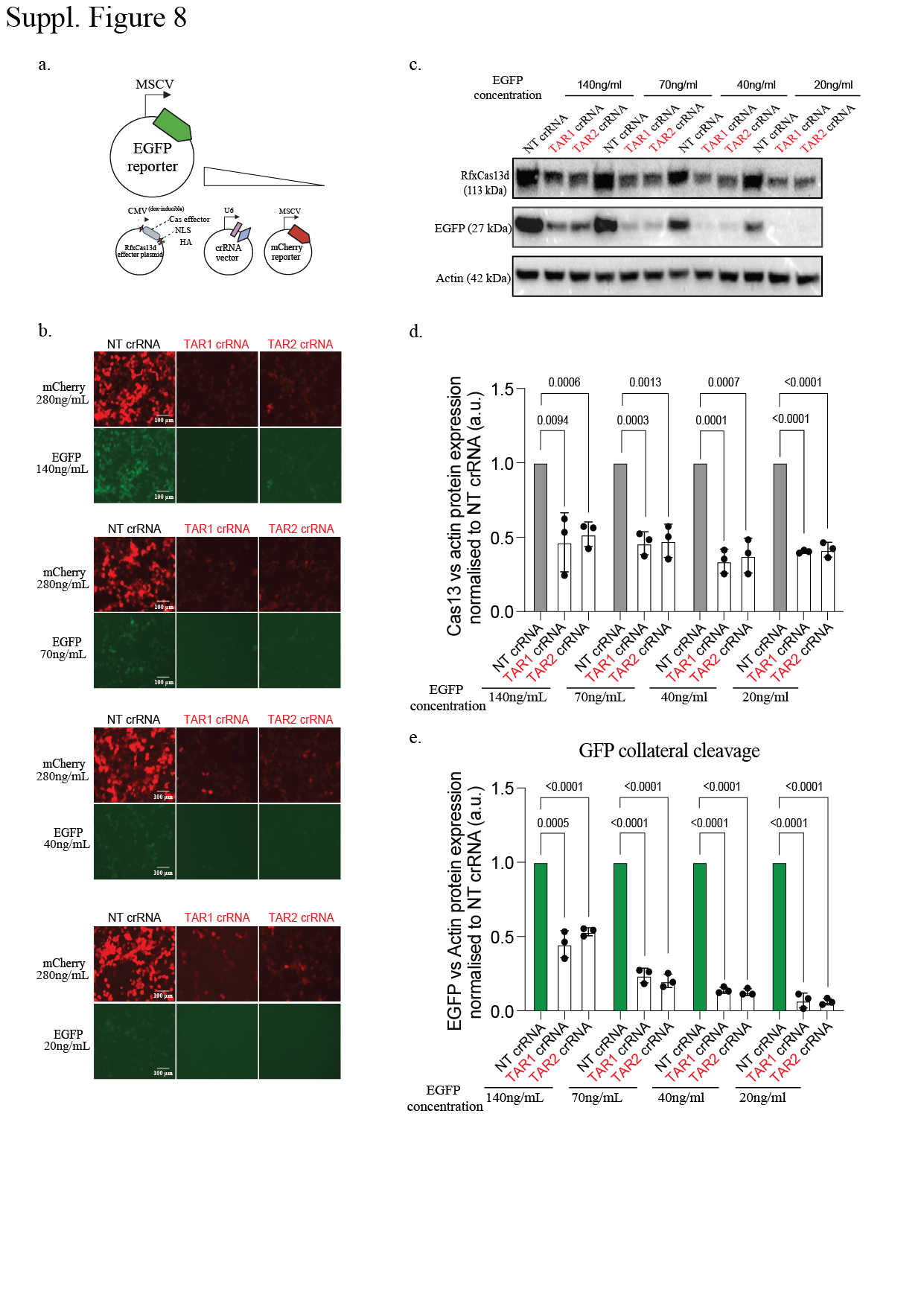

**Suppl. Figure 8**

**a,** Schematic of RfxCas13d on-target (mCherry) and collateral cleavage (EGFP) silencing with the titration of EGFP non-target abundance while maintaining mCherry target abundance. b, Representative fluorescence microscopy images of the silencing of mCherry and EGFP transcripts by RfxCas13d with two potent mCherry-targeting crRNAs versus a non-targeting (NT) control crRNA in HEK 293T cells at the highest EGFP concentration (280 ng/mL) and the lowest EGFP concentration (20 ng/mL). Scale bar is 100μm. **c,** Representative Western blot of the expression level of RfxCas13d, EGFP, and β actin protein levels in HEK293T cells expressing either mCherry-targeting crRNA or NT crRNA. **d-e,** Quantification of RfxCas13d and EGFP protein silencing normalised to β actin protein levels (n = 3 for each concentration). The data are represented in arbitrary units (a.u.). Errors are the s.e.m. and p values of Student’s t-test are indicated (95% confidence interval).

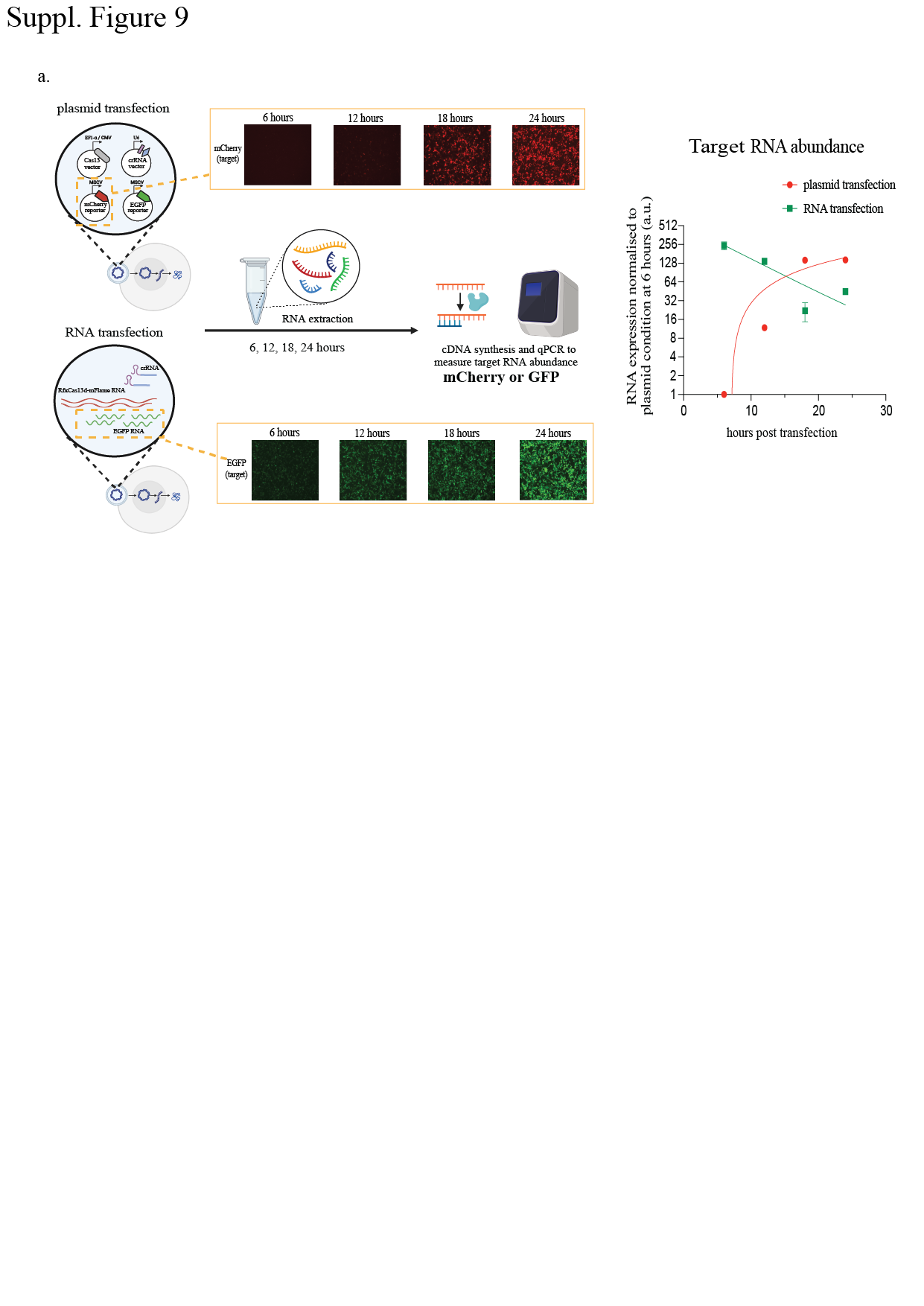
**Suppl. Figure 9**

**a,** **Left,** Schematic of the pipeline of RNA extraction from plasmid and RNA transfection assays at 6, 12, 18 and 24 hours post-transfection to assess target RNA abundance. **Right,** Quantification of relative target RNA expression levels for plasmid transfection (green) and RNA transfection (red) conditions at 6, 12, 18 and 24 hours post-transfection measured by qPCR.

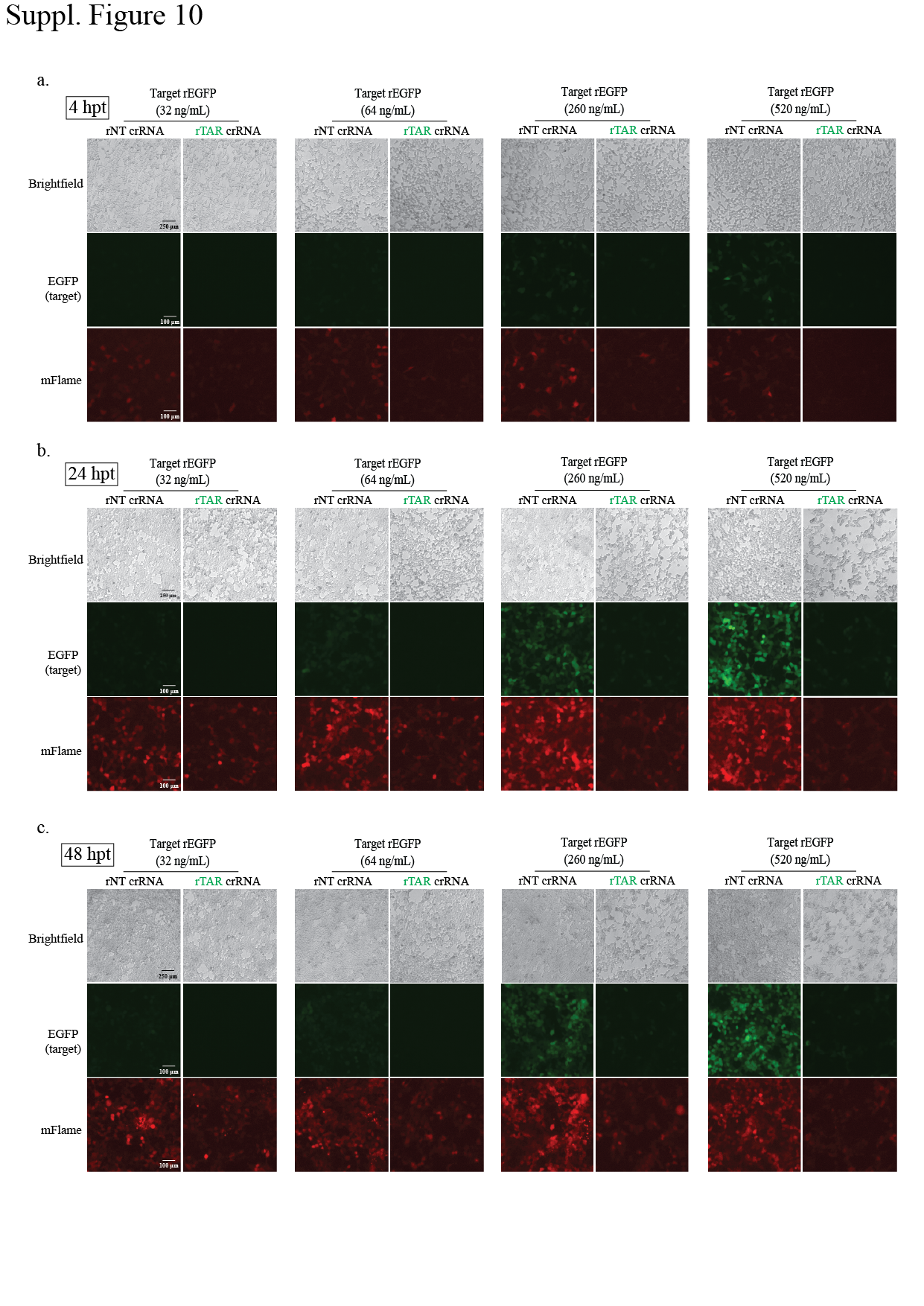

**Suppl. Figure 10**

**a-c,** Representative brightfield and fluorescent images of HEK293T cells at 4, 24, and 48 hours post-transfection of rRfxCas13d-mFlame, rEGFP, and either rNT or rEGFP-targeting crRNA. Target EGFP RNA is transfected at a titration of different concentrations ranging from 32ng/mL, 64ng/mL, 260ng/mL, and 520ng/mL (the concentrations used for proteomic analyses). Scale bar is 250μm for Brightfield images and 100μm for fluorescent images.

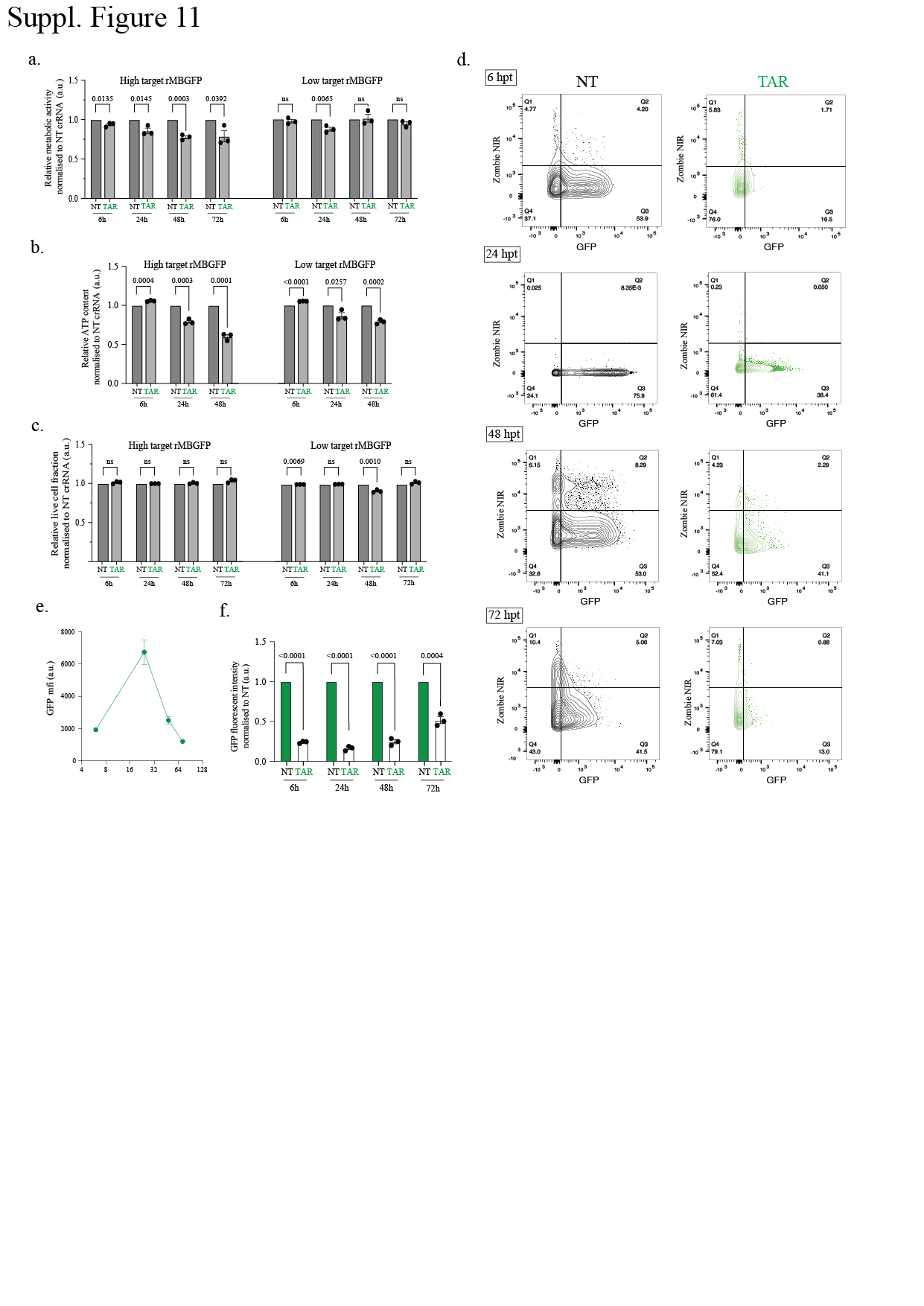
**Suppl. Figure 11**

**a-c,** Resazurin cell metabolism assay (quantification of cell metabolic activities), CellTiter-Glo (CTG) assay (quantification of metabolically active cells) and Zombie Near Infrared (ZIR) cell viability assay (quantification of percentage of cells alive) performed for HEK293T cells transfected with rEGFP, rRfxCas13d, rEGFP-targeting or non-targeting (NT) crRNA. Data points in the graph are averages of the normalised to the NT crRNA conditions for each time point (n = 3). Error bars are the s.e.m.. **d,** FACS plot of ZIR assay showing representative images of the silencing of rEGFP (high target abundance) as well as dead cell positively stained with ZIR dye. **e,** Quantification of EGFP mean fluorescence signal (a.u.) measured from Flow Cytometry in the NT crRNA condition (high target abundance) at different time points. **f,** Quantification of rEGFP silencing quantified from FACS data. Data points in the graph are averages of the normalised to the NT crRNA conditions for each Cas13 effector protein (n = 3).

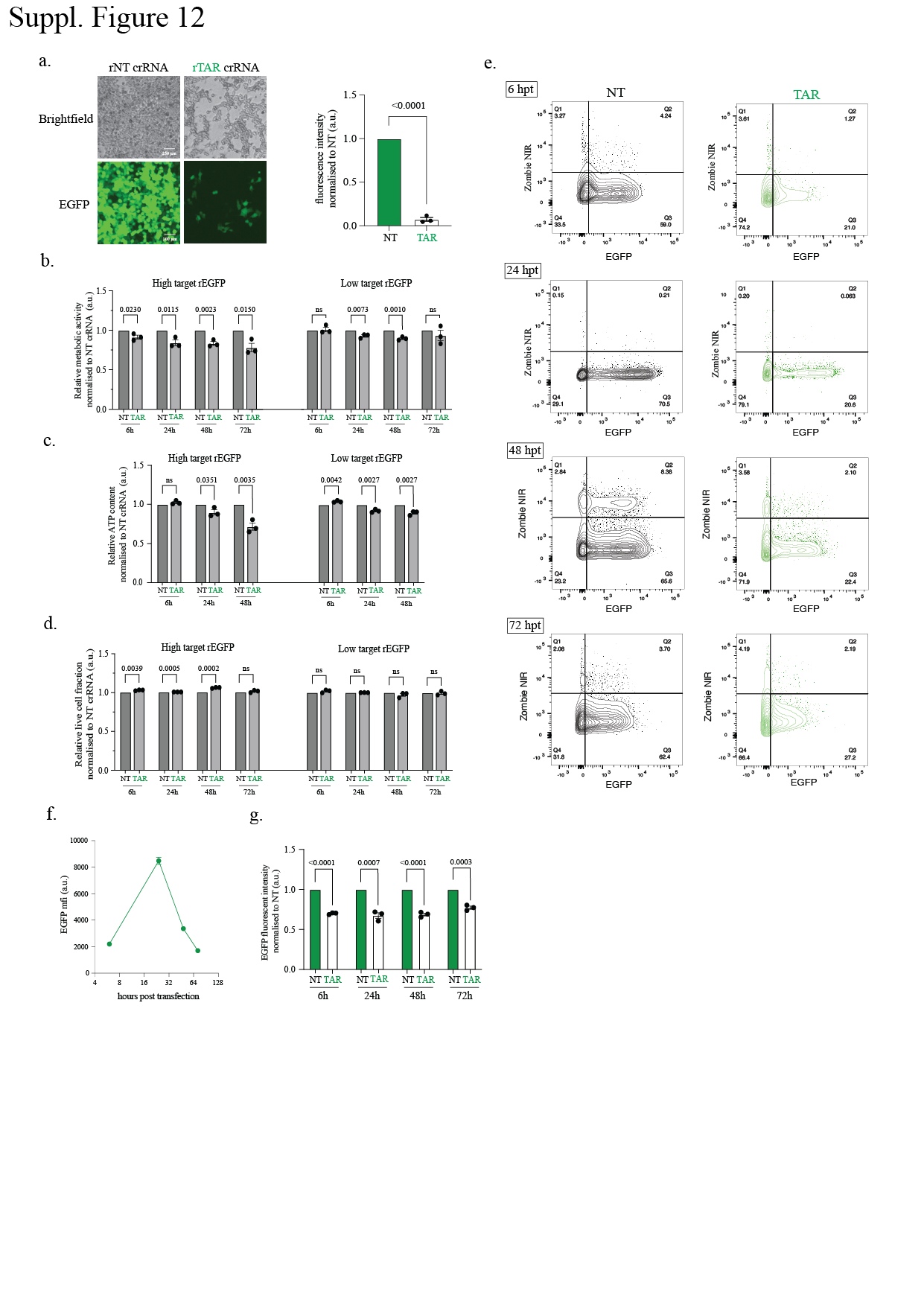

**Suppl. Figure 12**

**a, Left,** Representative brightfield and fluorescent images of HEK293T cells stably expressing RfxCas13d at 48h post transfection of rEGFP-targeting crRNA or rNT crRNA and rEGFP in the RNA form. Scale bar is 250μm for Brightfield images and 100μm for fluorescent images. **Right,** Quantification of RfxCas13d silencing of EGFP transcripts with either rNT or rEGFP-targeting crRNAs. Data points in the graph are averages of the normalised mean fluorescence from four representative fields of view imaged (n = 3). Errors are the s.e.m. and p values of Student’s t-test are indicated (95% confidence interval). **b-d,** Resazurin cell metabolism assay (quantification of cell metabolic activities), CellTiter-Glo (CTG) assay (quantification of metabolically active cells) and Zombie Near Infrared (ZIR) cell viability assay (quantification of percentage of cells alive) performed for HEK293T cells stably expressing RfxCas13d transfected with rEGFP, rEGFP-targeting or non-targeting (NT) crRNA at different time points. Data points in the graph are averages of the normalized to the NT crRNA conditions for each time point (n = 3). Error bars are the s.e.m.. **e,** **Left,** FACS plot of ZIR assay showing representative images of the silencing of rEGFP (high target abundance) as well as dead cell positively stained with ZIR dye. **f,** Quantification of EGFP mean fluorescence signal (a.u.) measured from Flow Cytometry in the NT crRNA condition (high target abundance) at different time points. **g,** Quantification of rEGFP silencing quantified from FACS data. Data points in the graph are averages of the normalized to the NT crRNA conditions for each Cas13 effector protein (n = 3).

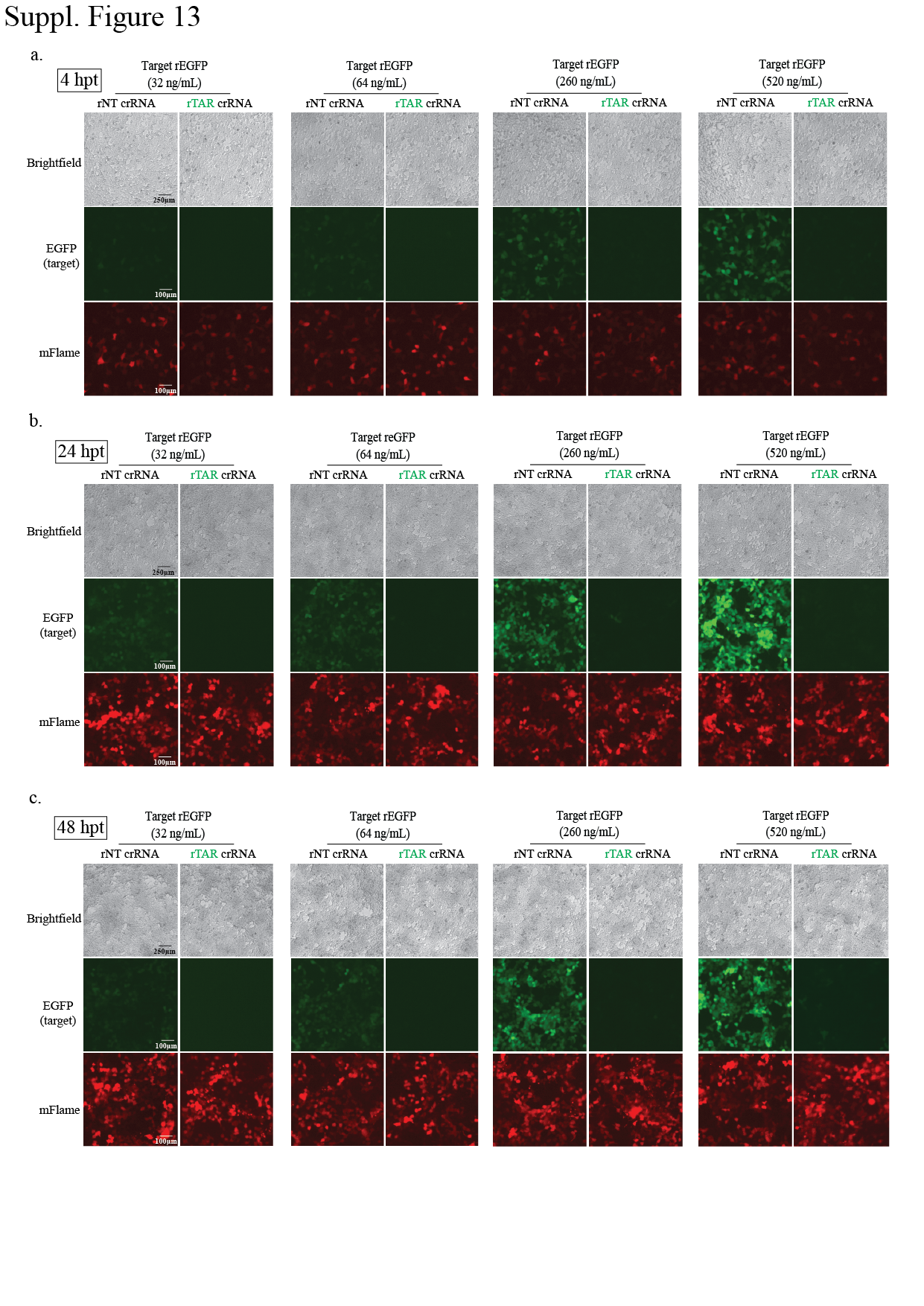
**Suppl. Figure 13**

**a-c,** Representative brightfield and fluorescent images of HEK293T cells at 4, 24, and 48 hours post-transfection of rPspCas13b-mFlame, rGFP, and either rNT or rGFP-targeting crRNA. Target EGFP RNA is transfected at a titration of different concentrations ranging from 32ng/mL, 64ng/mL, 260ng/mL, and 520ng/mL (the concentrations used for proteomic analyses). Scale bar is 250μm for brightfield images and 100μm for fluorescent images.

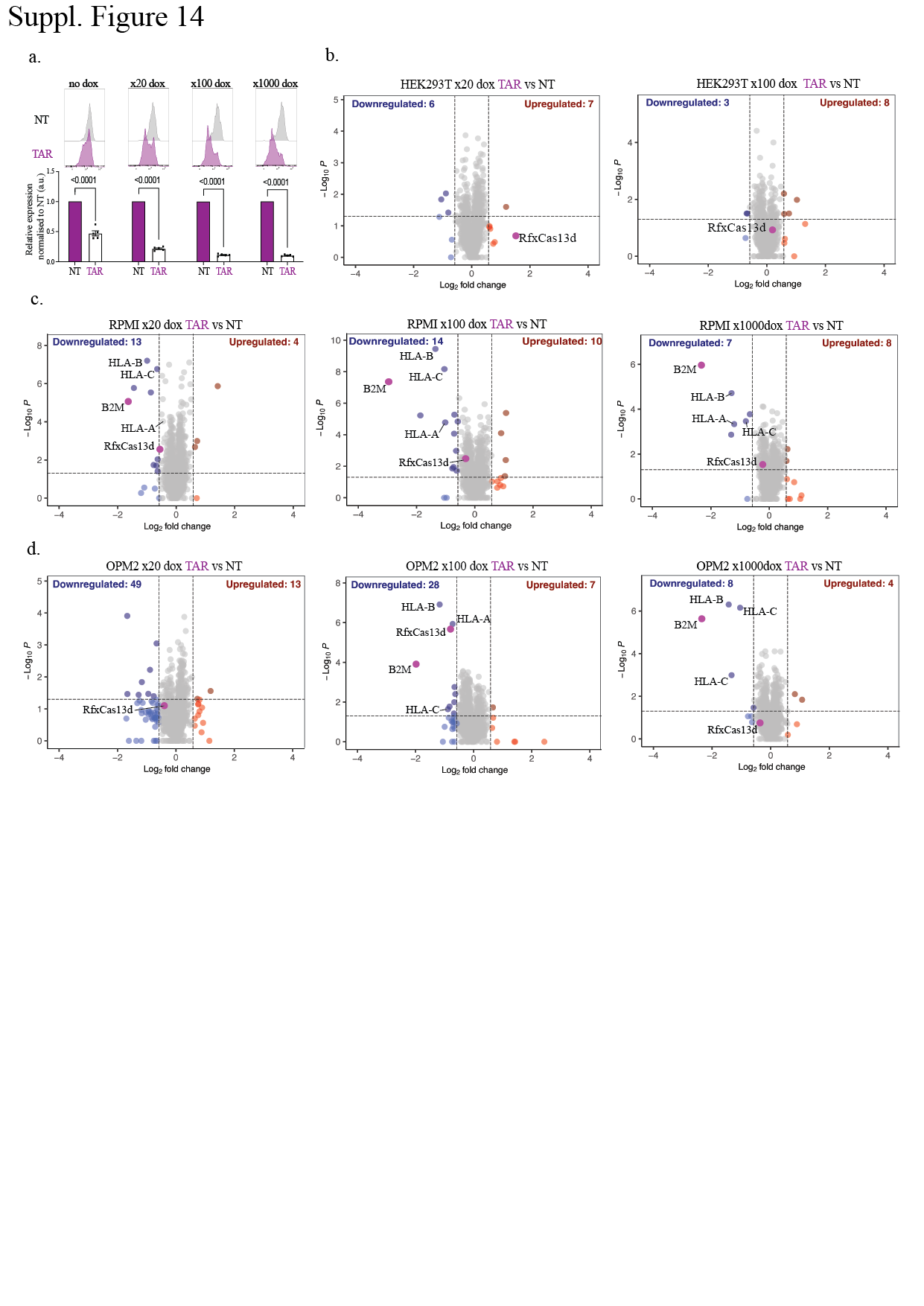

**Suppl. Figure 14**

**a,** Quantification of median β2m expression levels of cells stably expressing RfxCas13d and β2m-targeting or NT crRNA in HEK293T cells at 4 different dox induction concentration (1000ng/mL, 100ng/mL, 20ng/mL, 0ng/mL). Data points in the graph are averages of the normalised median PE-Cy7 FSC level (anti-β2m PE-Cy7 antibody) from four to six representative fields of view imaged (n = 3 or 5). The data are represented in arbitrary units (a.u.). Errors are the s.e.m. and p values of Student’s t-test are indicated (95% confidence interval). **b,** Volcano plots showing the proteome of cells stably expressing RfxCas13d and β2m-targeting or NT crRNA in HEK293T cells at two different dox activation concentration (100ng/mL, 20ng/mL) 48 hours post-activation. Data points represent proteins, with log_2_FC > 0.58 (upregulation, blue) or log_2_FC < −0.58 (downregulation, red) and p < 0.05 indicating significantly differential expression. Total downregulated and upregulated proteins are shown at the top corners. Cas13 and β2m are labelled purple. Results were analysed by an unpaired two-tailed Student’s t-test (n = 5). **c&d,** Volcano plots showing the proteome of cells stably expressing RfxCas13d and β2m-targeting or NT crRNA in RPMI and OPM2 cells at three different dox activation concentration (1000ng/mL, 100ng/mL, 20ng/mL) 48 hours post-activation. Data points represent proteins, with log_2_FC > 0.58 (upregulation, blue) or log_2_FC < −0.58 (downregulation, red) and p < 0.05 indicating significantly differential expression. Total downregulated and upregulated proteins are shown at the top corners. RfxCas13d and β2m are labelled purple. Results were analysed by an unpaired two-tailed Student’s t-test (n = 5).

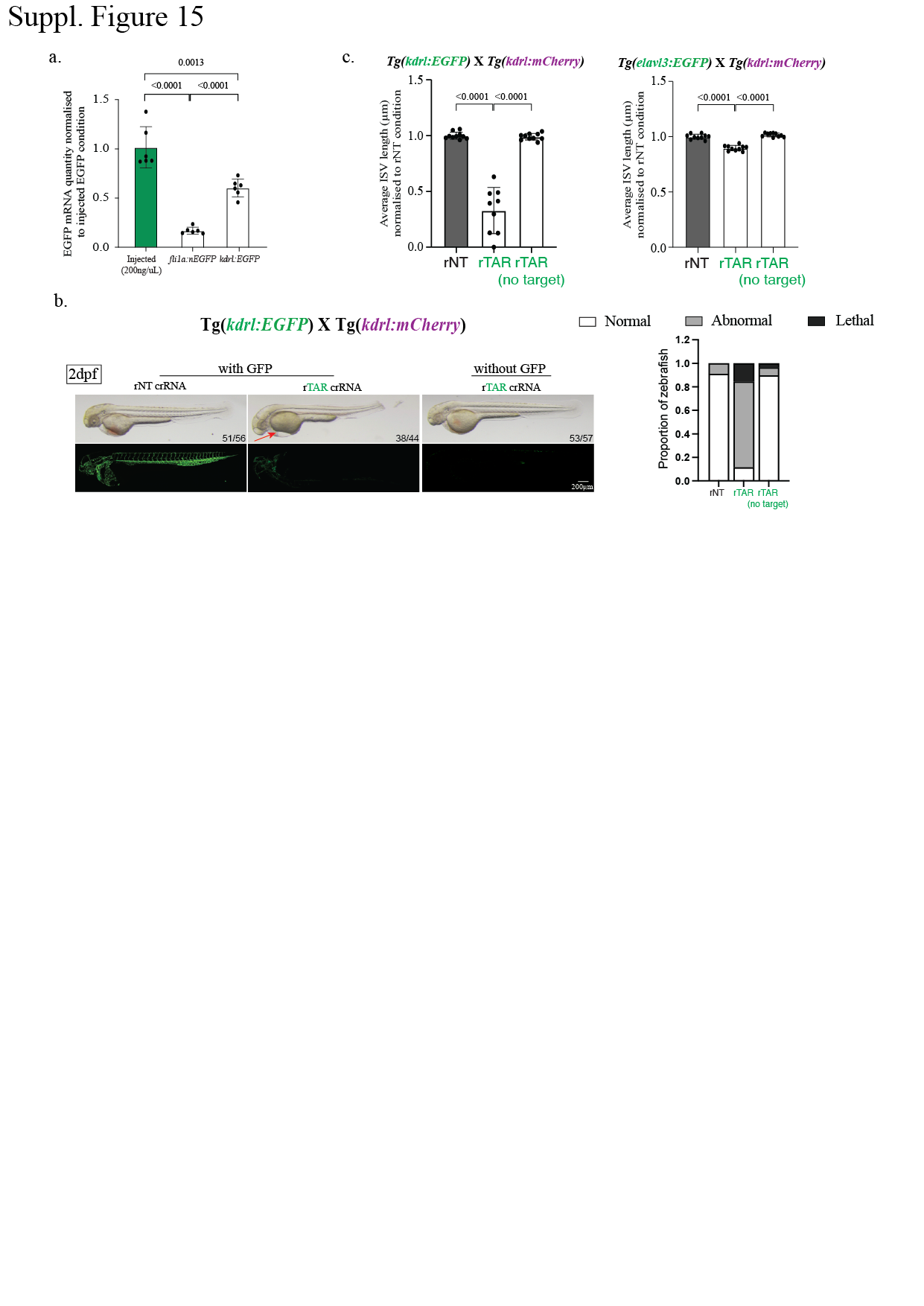
**Suppl. Figure 15**

**a,** Quantification of the expression of EGFP RNA measured by qPCR in *Tg(fli1a:EGFP)* and *Tg(kdrl:EGFP)* transgenics compared to zebrafish exogenously injected with EGFP at 200ng/μL. **b**, **Left**, Brightfield and fluorescent images of *Tg(kdrl:EGFP)*X*Tg(kdrl:mCherry)* double transgenic embryos from Fig. 5f showing embryo morphology as well as EGFP on-target cleavage using RfxCas13d with EGFP-targeting crRNA or NT crRNA control at 2 dpf. Non-transgenic siblings were also injected with RfxCas13d with EGFP-targeting crRNA as a negative control and displayed no morphological defects. Arrows indicate pericardial oedema. Numbers on the bottom right corner shows representative zebrafish out of all fertilised and survived zebrafish embryos. Scale bar is 200μm. **Right**, Quantification of the proportion of zebrafish that are normal (white), abnormal (grey) and lethal (black) after injection at 2dpf for *Tg(kdrl:EGFP)*X*Tg(kdrl:mCherry).* **c**, Quantification of average ISV length normalised to rNT crRNA negative control condition for *Tg(kdrl:EGFP)*X*Tg(kdrl:mCherry)* and *Tg(elavl3:EGFP)*X*Tg(kdrl:mCherry)*
